# Alternative cleavage of a bone morphogenetic protein (BMP) produces ligands with distinct developmental functions and receptor preference

**DOI:** 10.1101/131276

**Authors:** Edward N. Anderson, Kristi A. Wharton

**Author notes:** To whom correspondence should be addressed: Kristi A. Wharton.

## Abstract

TGF-β and Bone Morphogenetic Protein (BMP) family proteins are made as proprotein dimers, which are cleaved by proprotein convertases to release the active C-terminal ligand dimer. Multiple proteolytic processing sites in Glass bottom boat (Gbb), the *Drosophila* BMP7 ortholog, can produce distinct forms of active ligand. Cleavage at the S1 or atypical S0 site produces Gbb15, the conventional small BMP ligand, while cleavage at the NS site produces the larger Gbb38 ligand (1, 2). Here, we found that blocking NS cleavage increased association of the full length prodomain with Gbb15 resulting in a concomitant decrease in signaling activity. NS cleavage is required *in vivo* for Gbb-Decapentaplegic (Dpp) heterodimer-mediated wing vein patterning but not in cell culture to enable Gbb15-Dpp heterodimer activity. Gbb NS cleavage is also required *in vivo* for the regulation of pupal ecdysis and viability that is dependent on the type II receptor Wishful thinking (Wit). We found that the ability of Gbb38 to signal requires the expression of either Wit or the type I receptor, Saxophone (Sax). Finally, we discovered that the production of Gbb38 in 3rd instar larvae results when processing at the S1/S0 site is blocked by O-linked glycosylation. Our findings demonstrate that BMP prodomain cleavage can ensure that the mature ligand is not inhibited by the prodomain. Furthermore, alternative processing of BMP proproteins produces ligand types that signal preferentially through different receptors and exhibit specific developmental functions.

---

Bone Morphogenetic Proteins (BMPs), members of the TGF-β family of signaling proteins, have numerous developmental and physiological roles (3–5). Like other TGF-β family members, BMPs are synthesized as large 400-500 amino acid proproteins that form dimers linked by a C-terminal disulfide (6). The 110-140 amino acid ligand domain is proteolytically cleaved from the C-terminus by a proprotein convertase (PC) such as Furin. After secretion, the C-terminally derived ligand dimer binds and activates a complex of type I and type II transmembrane serine/threonine kinase receptors. In the active complex, the type II receptor phosphorylates the type I receptor that in turn phosphorylates downstream receptor-mediated Smad (R-Smad) signal transducers that act as transcription factors. While the vast majority of research in the field has focused on the activity of the ligand, there is a growing appreciation of the regulatory functions of BMP and TGF-β family prodomains (6–8). In general, it has been proposed that prodomains are important for proper folding, dimerization, and secretion of the mature ligand. A comparison of prodomains between different members of the TGF-β/BMP family shows a lower sequence conservation, in contrast to the high sequence conservation observed between their ligand domains. The low degree of prodomain sequence conservation might mean that TGF-β/BMP prodomains function as mere chaperones, as has been shown for prodomains of other proteins (9). However, this divergence in TGF-β/BMP prodomain sequences across the family could instead contribute to the functional diversification of signaling exemplified by TGF-β/BMP ligands (6).

The best studied example of a TGF-β family prodomain that plays an important role in ligand activity is TGFβ1. The TGFβ1 prodomain remains non-covalently associated with the ligand dimer after secretion, producing a latent complex that is inactive until the prodomain is removed from the ligand. Receptor activation is physically blocked by the prodomain which wraps around the mature TGFβ1 ligand domain, blocking receptor binding surfaces (10). Like TGFβ1, the non-covalent GDF8 prodomain-ligand complex is latent and prevents ligand-receptor binding (11). BMP ligands also form non-covalent complexes with their prodomains, but the consequences of these interactions are varied and less well understood. In most cases, BMP prodomain-ligand complexes are not latent (12) and ligand activity may instead be regulated by the prodomain in more subtle ways. In the case of BMP4, cleavage at a second PC site, S2, reduces prodomain-ligand association and is required for long-range signaling in *Xenopus* embryos (13, 14). During mouse development, cleavage at the BMP4 S2 site has been shown to be required for the development of specific tissues (15, 16).

In contrast, the BMP7 prodomain competes with the ligand for binding type II receptors *in vitro*, but this competition does not reduce signaling activity in the context of cell culture assays (17). Similarly, the BMP9 prodomain has been proposed to interact with type II receptors and to confer receptor specificity by blocking ligand binding to ACVR2A but not its binding to ACVR2B or BMPR2 type II receptors (18). In a separate study using real time surface plasmon resonance, the BMP9 prodomain was instead found to be rapidly displaced from the mature ligand by all receptor types, and in this case the prodomain did not cause latency (19). The variable findings regarding the role of BMP prodomains in receptor competition could reflect differences between the experimental approaches, or the context in which prodomain-ligand or prodomain-receptor interactions were examined. Therefore, studying the consequences of prodomain-ligand interactions *in vivo* is of particular interest.

Our laboratory has previously reported that BMP signaling activity can be affected by processing of the Gbb proprotein at different PC cleavage sites (1). In addition to the conventional S1 site that separates the conserved ligand domain from the more divergent prodomain, the *Drosophila* BMP7 ortholog Glass bottom boat (Gbb) contains an additional PC cleavage site, NS, within the N-terminal half of the prodomain (1, 2). Cleavage at the NS site produces a large ligand, Gbb38, whereas cleavage at the C-terminal S1 site produces the conserved smaller ligand, Gbb15. In many tissues, Gbb38 is more abundant than Gbb15 (1). In the developing wing epithelium, the NS site is required for wild-type signaling activity and range, while the S1 site is largely dispensable. Furthermore, the NS cleavage site has been shown to be necessary and sufficient for the rescue of *gbb* null lethality (2). In cell culture, both the NS and S1 cleavage sites have been shown to be critical to achieve full BMP signaling activity (1). Together, these findings indicate that Gbb38 is an active ligand with functions distinct from Gbb15. However, it is not exactly clear how cleavage influences the maturation of either ligand, or whether the prodomain impacts ligand activity and/or influences receptor preference. Furthermore, the suggestion that context-specific requirements for Gbb could be explained by the activities of different ligands produced by alternative processing requires further investigation.

The *gbb* gene is known to be required for multiple developmental and physiological processes in Drosophila. The roles of *gbb* in pattern formation have been shown to be mediated by receptor complexes composed of the type II receptor Punt, and type I receptors Thickveins (Tkv) and/or Saxophone (Sax), which phosphorylate the downstream R-Smad signal transducer Mothers against decapentaplegic (Mad) (20, 21). During wing development, both *gbb* and the BMP2/4 ortholog *decapentaplegic* (*dpp*) are required for wing patterning and differentiation (21, 22), with both homodimers and Gbb/Dpp heterodimers likely contributing to wing patterning (22–24). *gbb* and *dpp* loss of function mutations exhibit different effects on BMP signaling and produce distinct wing phenotypes, indicating that Gbb and Dpp are not functioning as an obligate heterodimer during larval wing patterning. However, in the case of the development of the posterior cross vein during pupal wing development, Gbb/Dpp heterodimers comprise the most likely active ligand (24–26). In the nervous system, Gbb signaling specifically requires the type II receptor Wit (27, 28). *wit* and *gbb* function is also required for the expression of peptide hormones that regulate pupal ecdysis behaviors (29). The varied roles for Gbb — cell fate specification, synapse growth promotion, regulation of neurotransmission and hormone expression — could be produced by a mechanism that regulates specific outcomes of Gbb signaling, potentially by allowing activation of different sets of receptors. Given the differences in Gbb38 and Gbb15 accumulation between tissues (1), we considered the possibility that the Gbb prodomain is involved not only in regulating the production of specific ligand forms but also their signaling output.

We examined in detail the processing of proGbb, prodomain-ligand interactions, and the *in vivo* requirements for NS cleavage. We found that S2 cells secrete Gbb15 in complex with the NS-cleaved prodomain. Blocking NS cleavage reduced Gbb15 signaling activity, and increased the association of Gbb15 and the uncleaved prodomain. Signaling activity by Gbb38 was dependent on Sax or Wit. *In vivo*, we found NS cleavage is required for wing vein patterning and *wit*-dependent pupal ecdysis. In 3rd instar larvae, O-linked glycosylation blocks S1 cleavage, and Gbb38 is produced as a ligand that preferentially activates Wit. Overall, our results demonstrate that regulated alternative cleavage of the Gbb prodomain produces ligands that preferentially activate specific receptors.

## RESULTS

*Gbb prodomain cleavage products are secreted*—To understand how NS cleavage in the Gbb prodomain may impact signaling, we first wanted to identify all cleavage products, their ability to be secreted, and how they were affected by mutations in specific PC sites. We raised antibodies against specific prodomain epitopes, and with the existing C-terminal directed α-GbbC (1), we were able to identify all possible products cleaved from the proGbb precursor protein (Fig. 1A). α-GbbN recognizes amino acids (aa) 46-61, near the N-terminus of the prodomain, and was used to identify N-terminal cleavage products. α-GbbCore, named after the Core/Arm domain of BMP2 and TGFβ1 (10, 30), recognizes aa 127-144 and any cleavage products resulting from the removal of the N- or C-terminus. When wild type *gbb* was expressed by *Drosophila* Schneider 2 (S2) cells, we detected secreted Gbb prodomain cleavage products in the media (Fig. 1B). α-GbbN identified a 10 kDa band that matched the expected size of the NH_3_-NS fragment on Western blots of the conditioned media. α-GbbCore detected a 28 kDa fragment which matched the expected size of the NS-S1 cleavage product, and a previously unidentified prodomain fragment (31). Finally, α - GbbC detected the expected Gbb15 ligand secreted into the media (Fig. 1C). In cell lysates, we found high levels of proGbb, and detected Gbb38 and other prodomain cleavage products at lower abundance (Fig. S2A). Taken together, we found that in S2 cells the Gbb propeptide is cleaved at both the NS and S1 sites, and that all resulting cleavage products are secreted.

**Figure 1:**
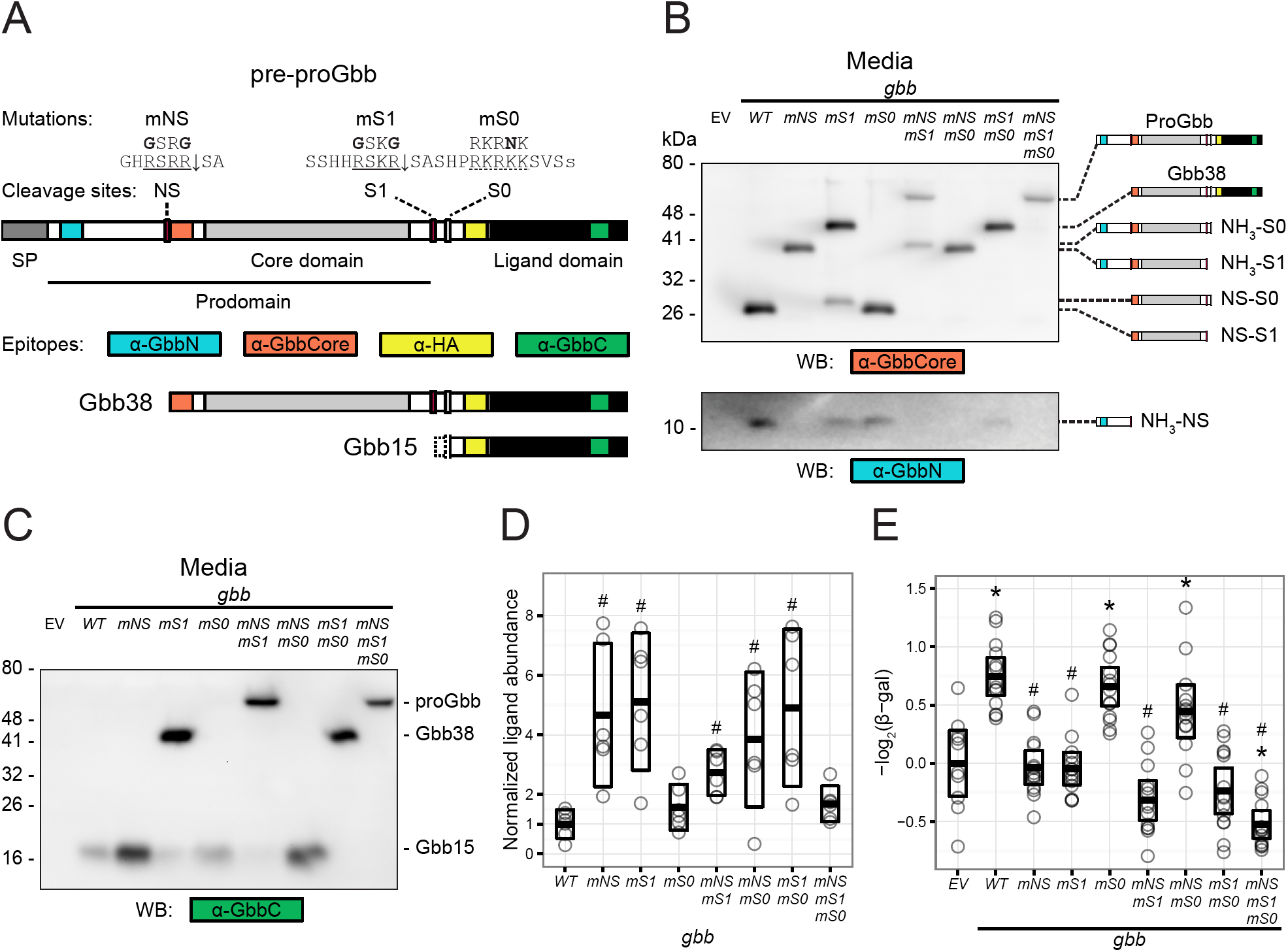
Gbb prodomain cleavage affects ligand abundance and activity. A, Gbb protein schematic, showing domains, signal peptide (SP), cleavage sites, mutations, and location of antibody epitopes. Furin cleavage motifs at NS and S1 are underlined, with the downward arrow indicating the cleaved bond. The atypical S0 cleavage site is underlined with a dashed line. B, C, reducing Western blots of Gbb cleavage products secreted by S2 cells expressing *gbb* cleavage mutants, using antibodies that recognize specific epitopes in the Gbb prodomain (B) or the C-terminal ligand domain (C). D, quantification of secreted ligand abundance for representative Western blot shown in (C). For mutant constructs that produce multiple forms of secreted Gbb ligand, the summed total of both forms is shown. E, steady-state signaling activity produced by gbb cleavage site mutant constructs when co-expressed with the *BrkSE-LacZ* reporter in S2 cells. Expression of *BrkSE-LacZ* reporter is repressed by BMP signaling, and is shown here as opposite log transformed β-gal activity, with the activity of empty vector (EV) transfected cells set to zero. D, E, Bars indicate the mean and 95% confidence intervals (CI). * indicates *p* < 0.05 compared to EV, # indicates *p* < 0.05 compared to WT *gbb*, using the general linear hypothesis test (GLHT) for multiple comparison testing.

We next generated *gbb* constructs that harbor all combinations of mutations in the PC sites: NS, S1, and the atypical S0 (referred to as the "Shadow" site by (2)). When NS cleavage is blocked (*mNS*), the NH_3_-NS product is absent, a 37 kDa band corresponding to the intact prodomain (NH_3_-S1) is detected, and an approximately 5-fold increase in Gbb15 is observed (95% confidence limits (CL) 1.7-13.3 fold, Fig. 1D). When S1 cleavage is blocked (*mS1*), the most abundant product is Gbb38, and the NH_3_-NS fragment is also detected in the media (Fig. 1B-D). The appearance of NS-S0 and the low abundance of Gbb15, when compared to total Gbb ligands (22%, 95% CL 9%-35%, Fig. S2A), indicate that when S1 is mutated, some cleavage most likely occurs at the atypical S0 site (Fig. 1B, C). Blocking S0 cleavage (*mS0*) or S0 in combination with NS (*mNSmS0*) had no detectable effect on the presence or abundance of prodomain cleavage products or Gbb15 (Fig. 1B-D). However, blocking both S1 and S0 (*mS1mS0*) eliminated the NS-S0 fragment present in mS1 and resulted in the secretion of Gbb38 at much higher abundance than Gbb15 produced by a wild type construct (4.8-fold, 95% CL 1.7-13.5 fold, Fig. 1B-D). When cleavage at both NS and S1 is blocked (*mNSmS1*), the entire uncleaved proGbb is detected in the media, along with NH_3_-S0 and very low levels of S0-cleaved Gbb15 (Fig. 1B-D). Finally, mutating all three PC sites (*mNSmS1mS0*) resulted in a loss of all cleavage products and the secretion of proGbb (Fig. 1B, C).

To identify the putative active ligands that exist as dimers, we examined secreted Gbb products using a non-reducing Western blot. A Gbb15 dimer, detectable as a 40 kDa band, was present in the conditioned media of S2 cells expressing wild-type *gbb* (Fig. S2B). As we observed using reducing Western blots (Fig. 1), blocking NS cleavage (*mNS*) increased Gbb15 abundance, but blocking S0 cleavage (*mS0*) had little effect. Gbb38 dimers and proGbb dimers were secreted by cells expressing *gbb* ^*mS*1*mS0*^ and *gbb* ^*mNSmS1mS0*^, respectively. When S1 cleavage was blocked (*mS1*), we observed secretion of abundant Gbb38 dimers, as well as Gbb15-Gbb38 dimers where only one monomer in the dimer was cleaved at the S0 site. In this case, Gbb15 dimers produced by S0 cleavage of both monomers were present at much lower abundance. When cleavage at both NS and S1 was blocked (*mNSmS1*), a mix of proGbb dimers, Gbb15-proGbb, and low abundance Gbb15 dimers were secreted. These data provide further evidence of the inefficiency of cleavage of the S0 site in S2 cells. Since the S0 site does not have the minimal RXXR Furin cleavage site (32), it may instead be cleaved by another PC such as the *Drosophila* PC2 ortholog Amontillado, which does not have significant activity in S2 cells (33).

*S1 cleavage is sufficient for signaling activity in S2 cells*—We measured the signaling activity of *gbb* cleavage mutants in S2 cells, using the *BrkSE-LacZ* reporter. In this assay, constitutive expression of LacZ is transcriptionally silenced in response to the BMP signaling-dependent *brinker* silencer element (34). In cells transfected with wild type *gbb*, signaling activity calculated as log_2_ (*BrkSE-LacZ*) is increased compared to endogenous levels as seen in the control EV transfected cells (Fig. 1E). When cells are transfected with a construct with all cleavage sites blocked (*mNSmS1mS0*), signaling activity is lower than in EV transfected cells. This likely reflects the inhibition of endogenous signaling by the formation of inactive proGbb-heterodimers with endogenously expressed BMPs (35, 36), and is consistent with our finding that expression of *gbb* ^*mNSmS1*^ in wing discs led to a cell autonomous loss of endogenous pMad (1). The level of BMP signaling as measured by the *BrkSE-LacZ* assay is significantly reduced when NS or S1 cleavage is blocked (*mNS* or *mS1*) (Fig. 1E), indicating a requirement for cleavage at each site. Blocking S0 cleavage (*mS0*) had no effect on signaling activity, and no activity is detectable when only S0 cleavage is permitted (*mNSmS1*). Therefore, cleavage at the S0 site alone is neither necessary nor sufficient for signaling activity in S2 cells.

Interestingly, while cleavage at only the NS site has been shown to fully rescue the lethality associated with *gbb* null alleles *in vivo* (2), we did not detect signaling activity in S2 cells transfected with *gbb* ^*mS1mS0*^ using the *BrkSE-LacZ* assay. Even though Gbb38 is produced when proGbb is cleaved at only the NS site in cells expressing *gbb* ^*mS1mS0*^ (Fig. 1B), NS cleavage alone is not sufficient to produce signaling activity in S2 cells (Fig. 1E). However, when only S1 cleavage is permitted (*mNSmS0*), we observe signaling activity (Fig. 1E). Thus, in S2 cells we find that NS cleavage is necessary but not sufficient for signaling activity, and S1 cleavage is both necessary and sufficient. We considered the possibility that NS cleavage is necessary to ensure full Gbb15 activity.

*Gbb15 activity is inhibited by the prodomain when NS cleavage is blocked*—The *BrkSE-LacZ* assay readout reflects multiple aspects of ligand production, secretion, receptor interaction, and downstream feedback, all in the context of BMP ligands endogenously expressed in S2 cells. To eliminate contributions of other signaling mechanisms that could modulate Gbb activity, we directly measured the activity of the secreted forms of Gbb15 and Gbb38 and compared their signaling kinetics. Conditioned media was produced by S2 cells stably transfected with *gbb*, *gbb* ^*mNS*^, or *gbb* ^*mS1mS0*^ expression constructs or empty vector. Since the concentration of secreted ligands (Gbb15 or Gbb38) that result from cells expressing cleavage mutant constructs (*mNS* or *mS1mS0*) is much higher than Gbb15 produced by wild-type *gbb* (Fig. 1C, D), we adjusted the cleavage mutant conditioned media so that ligand concentrations were equivalent, using conditioned media from EV expressing cells.

We measured signaling as the phosphorylation of Mad in S2 cells transiently transfected with FLAG-tagged *Mad* (*Mad-FLAG*). After a two-hour treatment of *Mad-FLAG* expressing cells, wild-type conditioned media, which contains Gbb15 and the NH_3_-NS and NS-S1 prodomain fragments, induced high levels of phosphorylated Mad (pMad), relative to total Mad-FLAG (Fig. 2A). Conditioned media from *gbb* ^*mNS*^, containing Gbb15 and the uncleaved prodomain NH_3_-S1, produced pMad at 55% of WT levels (*p* < 0.001, 95% CI 38%-79%). Therefore, we conclude that the activity of Gbb15 is reduced in the presence of the uncleaved prodomain (NH_3_-S1) suggesting that a failure to cleave the prodomain at the NS site can antagonize Gbb15-induced signaling. We also found that conditioned media from *gbb* ^*mS1mS0*^, which contains Gbb38 and the NH_3_-NS fragment but no Gbb15, produced pMad at 17% of WT levels (*p* < 0.001, 95% CI 12%-25%) (Fig. 2A), indicating that in this experimental system Gbb38 can induce signaling in S2 cells, albeit at lower levels than Gbb15. The difference in Gbb38 activity between this assay and the *BrkSE-lacZ* assay could indicate that the overexpressed *gbb* ^*mS1mS0*^ is inhibiting the activity of endogenously expressed BMPs (35, 36), or that some feedback mechanism reduces the ability of S2 cells that are expressing *gbb* ^*mS1mS0*^ to receive a Gbb38 signal.

**Figure 2:**
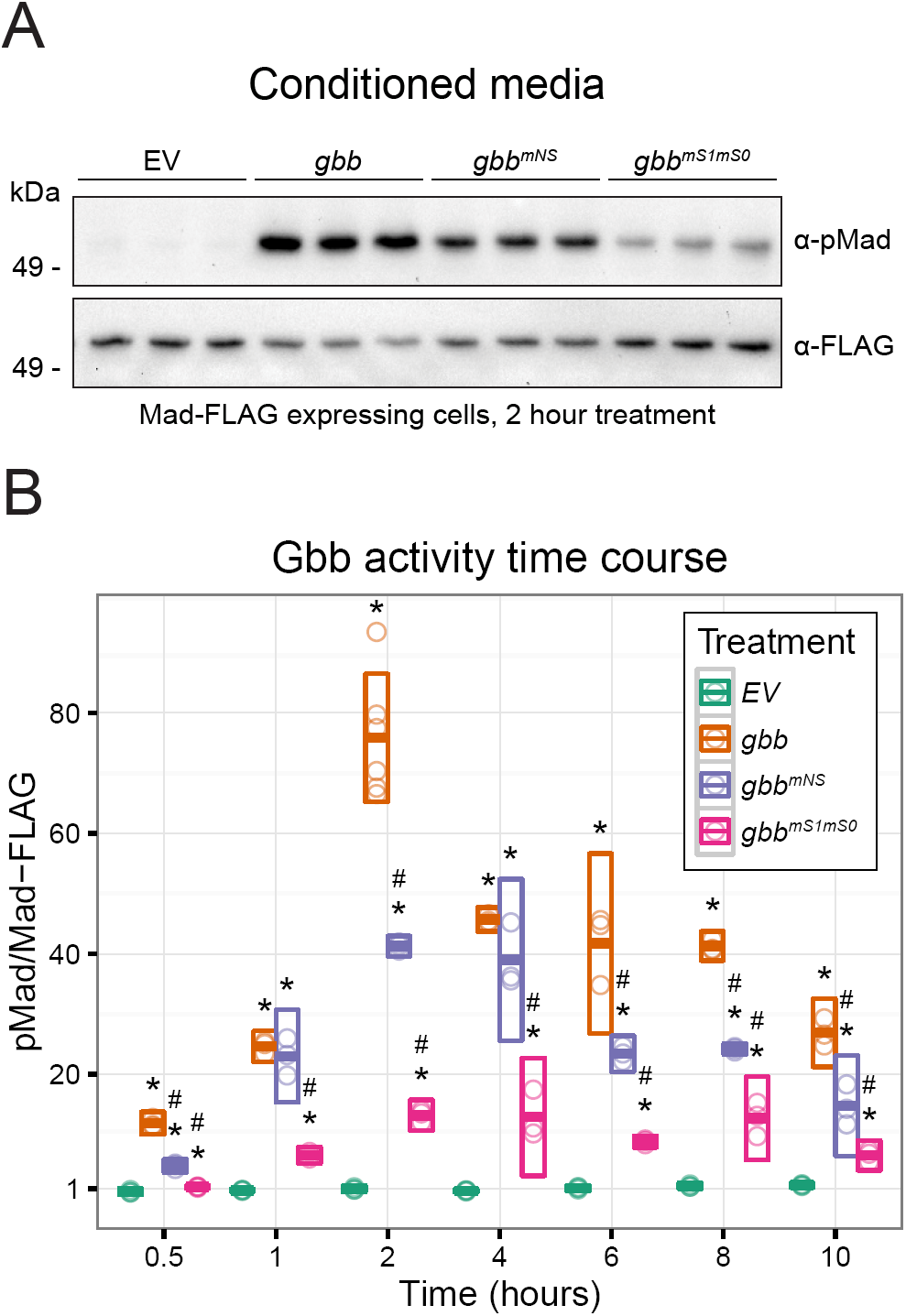
Activity and kinetics of Gbb cleavage mutant ligands. A, Western blots of phosphorylated Mad-FLAG (pMad) in lysates of S2 cells treated with *gbb* conditioned media for 2 hours. B, quantification of Mad-FLAG western blots shown in (A), also showing time course of *gbb* induced pMad from 0.5 to 10 hours. Bars indicate mean and 95% CI. At each time point, * indicates *p* < 0.05 compared to EV, # indicates p < 0.05 compared to WT *gbb*, using the GLHT multiple comparison test.

NS site cleavage of the Gbb ortholog, Scw, has been shown to be required for wild-type signaling kinetics (37). Unlike Gbb, Scw homodimers have very low signaling activity in S2 cells, and is instead thought to function primarily as a part of Scw-Dpp heterodimers (38). Scw-Dpp heterodimers show relatively rapid signaling kinetics, and exhibit peak activity at 1 hour (37). Mutating the Scw NS cleavage site was shown to alter heterodimer such that activity reached wild-type levels only after 3-5 hours, suggesting that a failure to cleave the Scw prodomain at the NS site impacted the signaling kinetics of Scw-Dpp heterodimers. Given the reduced level of signaling observed from *gbb* ^*mNS*^ conditioned media describe above, we also tested the effect of alternative processing of Gbb on signaling kinetics. We examined pMad levels produced over a time course of 0.5-10 hours in cells treated with conditioned media produced by different *gbb* constructs (Fig. 2). Activity of wild-type *gbb* conditioned media was detectable at 0.5 hours, peaked at 2 hours, and gradually decreased through hour 10. When NS cleavage was blocked, *gbb* ^*mNS*^ conditioned media had reduced activity at nearly all time points, with peak activity at 2-4 hours. When S1/S0 cleavage was blocked and only Gbb38 was present in conditioned media, we again detected low levels of signaling activity, with a peak at 2-4 hours. The low level of signaling at each time point was consistent with our finding that in S2 cells Gbb38 has less activity than Gbb15, though it signals with kinetics similar to Gbb15. In fact, preventing cleavage at NS or at S1/S0 did not affect the overall signaling kinetics. Given that blocking NS cleavage enabled secretion of the uncleaved prodomain with Gbb15, and that this led to a reduction in the ability of Gbb15 to signal (Fig. 1, 2A), results from the time course indicate that NS cleavage does not affect Gbb15 signaling kinetics.

*Cleavage at the NS site reduces Gbb15-prodomain association*—Cleavage of the GDF8/GDF11 prodomain by Tolloid metalloproteases, at a site that aligns near the NS site of Gbb, releases the ligand from the latent complex (39, 40). We next hypothesized that cleavage of the Gbb prodomain at the NS site may enable full levels of Gbb15 signaling by preventing prodomain-Gbb15 association. To examine prodomain-ligand interactions, Gbb with a C-terminal HA tag was co-immunoprecipitated from conditioned media using antibodies that recognize epitopes in each cleavage product. α-GbbN directly precipitates the NH_3_-NS fragment, and co-precipitates Gbb15 from conditioned media produced by cells expressing WT *gbb-HA* (Fig. 3). Similarly, α-GbbCore directly precipitates the NS-S1 fragment, and co-precipitates Gbb15. However, when Gbb15 is precipitated with α-HA, no co-precipitation of the NH_3_-NS or the NS-S1 cleavage product is observed. When NS cleavage is blocked, co-precipitation of Gbb15 using either α-GbbN or α-GbbCore to pull down the uncleaved prodomain is more efficient than with the NS cleaved prodomain, and α-HA effectively co-precipitates the NH_3_-S1 product. Finally, when S1/S0 cleavage is blocked the NH_3_-NS fragment co-precipitates with Gbb38 more efficiently than with Gbb15. Therefore, we conclude that Gbb15 forms a loosely associated complex with both NS-cleaved prodomain fragments (NH_3_-NS and NS-S1), and a failure to cleave at the NS site increases Gbb15-prodomain association. These co-immunoprecipitations were carried out at pH 6.5, the physiological pH of S2 cell culture media and larval hemolymph (41). Interestingly, at pH 7.4, we observed reduced co-precipitation of prodomain cleavage products with either Gbb15 or Gbb38, suggesting that these complexes are sensitive to pH (Fig. S3). We also observed pH-dependent interactions between Gbb ligands and prodomain cleavage products using heparin chromatography (Fig. S4). Together with the observation that blocking NS cleavage increases Gbb15 abundance but reduces its ability to signal (Fig. 1, 2), we propose that NS cleavage is required to prevent latency of Gbb15.

**Figure 3:**
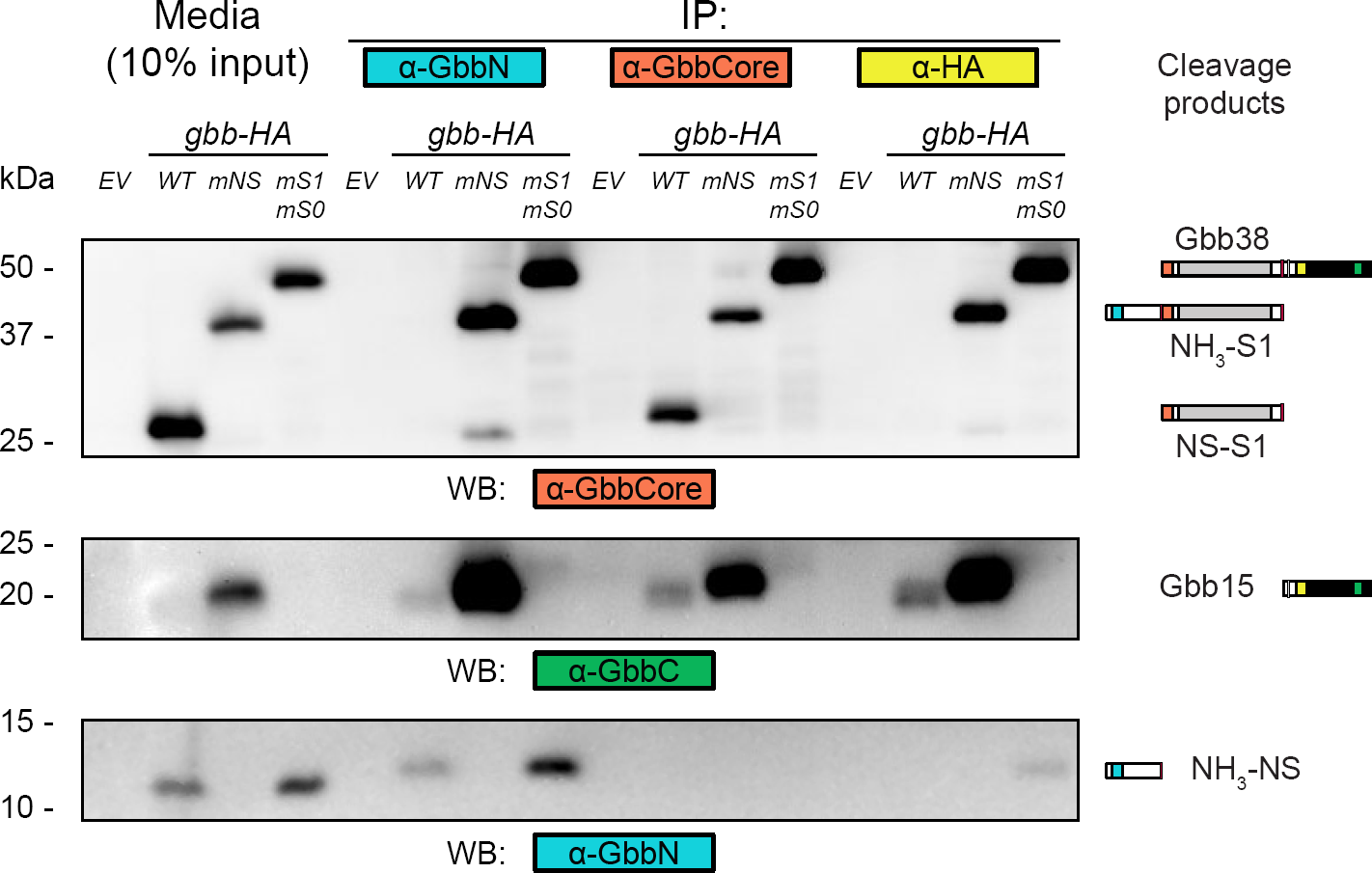
Cleavage of the NS site reduces Gbb15-prodomain association. Reducing Western blots of coimmunoprecipitating Gbb cleavage products. Conditioned media from S2 cells expressing *gbb-HA* cleavage mutants was immunoprecipitated with α-proGbb and α-CoreGbb to directly pull down prodomain cleavage products, and α-HA to pull down C-terminal cleavage products. Immunoprecipitate was analyzed by western blot using α-proGbb, α-CoreGbb, and α-GbbC to identify each cleavage product.

*Gbb alternative cleavage generates ligands that exhibit receptor preference*—Given that S2 cells express the type II receptor ACVR2A/B ortholog Punt but not the BMPR2 ortholog Wit (42), we considered the possibility that different behaviors of *mNS*, *mS1*, and *mS1mS0* in S2 cells, wing discs and in null rescues (1, 2) could reflect the usage of different receptors. To measure signaling activity in the context of different type II receptors, we co-transfected S2 cells with constructs expressing *punt* or *wit* and *Mad-FLAG*, and measured the level of pMad induced by conditioned media from cells expressing *gbb*. In cells transfected with *punt*, wild-type *gbb* conditioned media induced a 5-fold increase in pMad, compared to untreated cells (Fig. 4A). However, neither *gbb* ^*mNS*^ (NH_3_-S1 + Gbb15) nor *gbb* ^*mS1mS0*^ (NH_3_-NS + Gbb38) conditioned media induced a detectable increase in pMad over the empty vector conditioned media control. This failure to induce Mad phosphorylation suggests that, in the case of *gbb* ^*mNS*^ media, the ability of Gbb15 to induce Punt-mediated signaling is prevented when NS cleavage is blocked, and in the case of *gbb* ^*mS1mS0*^ media, Gbb38 cannot activate Punt-dependent signaling. In cells transfected with *wit*, conditioned media from all three *gbb*-expressing constructs elicited a significant increase in pMad with both wild-type *gbb* and *gbb* ^*mNS*^ conditioned media inducing an 8-fold increase, and *gbb* ^*mS1mS0*^ conditioned media resulting in a 5-fold increase in pMad. Taken together, these results indicate that activation of Punt by Gbb requires both NS and S1/S0 cleavage, and that neither Gbb38 associated with the NH_3_-NS fragment, nor Gbb15 with the uncleaved prodomain can activate Punt. On the other hand, both Gbb38 (*mS1mS0*) and Gbb15 with the uncleaved prodomain (*mNS*) can activate Wit-mediated signaling. Thus, ligands resulting from a mutation in either of the proconvertase processing sites can signal through the Wit type II receptor, and Gbb38 appears to function as a ligand that preferentially activates Wit.

**Figure 4:**
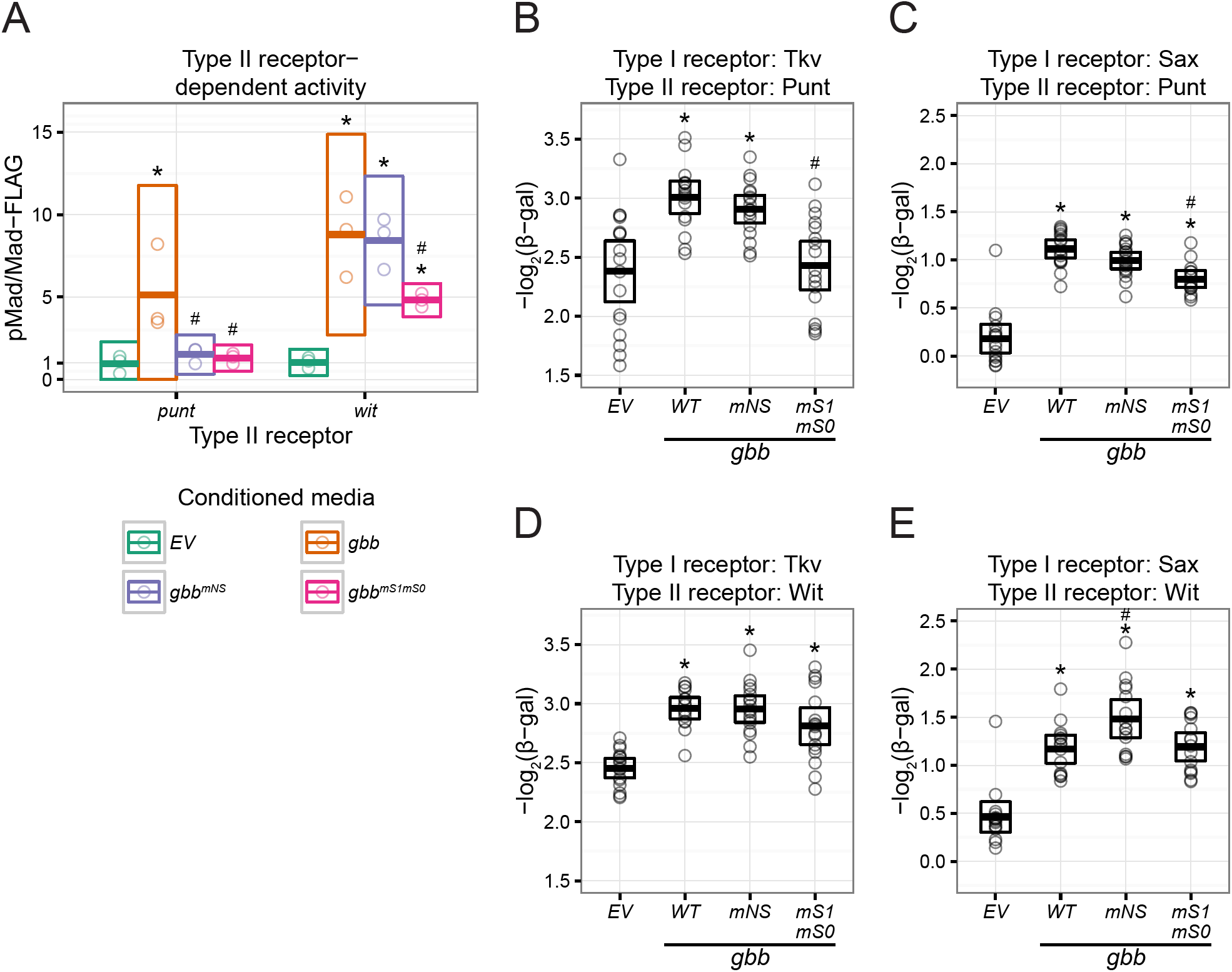
Receptor-specific activity of Gbb cleavage mutants. A, Type II receptor-dependent pMad induced by Gbb cleavage mutants. S2 cells expressing *Mad-FLAG* and *punt* or *wit* were treated with *gbb* conditioned media for 2 hours, and pMad was measured by western blot. B-E, steady state *BrkSE-LacZ* signaling assay of *gbb* co-expressed with all combinations of the type I receptors *tkv* or *sax*, and the type II receptors *punt* or *wit*. Bars indicate mean and 95% CI. * indicates *p* < 0.05 compared to EV, # indicates *p* < 0.05 compared to WT gbb, using GLHT multiple comparison test.

Gbb has been proposed to form high-affinity complexes with the type I receptor Sax, but not with Tkv (43). Consistent with this report, our laboratory previously found that Gbb38 co-immunoprecipitates with Sax but not Tkv (1). We further investigated the ability of different type I and type II receptor combinations to mediate signaling elicited by different Gbb cleavage products. *gbb* cleavage mutant constructs were co-expressed in S2 cells with constructs expressing type I receptors encoded by *tkv* or *sax* and type II receptors encoded by *punt* or *wit*, and signaling activity was measured using the *BrkSE-LacZ* assay. In cells expressing *tkv* and *punt*, we saw that blocking NS cleavage of Gbb did not reduce signaling activity (Fig. 4B). However, blocking S1/S0 cleavage (allowing production of only Gbb38) abolishes signaling activity. In cells expressing *sax* and *punt*, blocking NS cleavage, again, did not significantly reduce activity, but blocking S1/S0 cleavage (Gbb38) reduced but did not eliminate signaling (Fig. 4C). In cells expressing *tkv* and *wit*, mutations in either the NS or S1/S0 cleavage sites had no significant effect on signaling activity (Fig. 4D) consistent with our finding that Wit is refractory to the presence of the prodomain, either in association with Gbb15, or when the Core domain is included as an integral part of Gbb38. Like cells expressing *tkv* and *wit*, cells expressing *sax* and *wit* exhibit significant signaling from all three *gbb* constructs. In this context, *gbb* ^*mNS*^ produced an unexpected significant increase in activity over wild-type levels (Fig. 4E). Overall, we conclude that Gbb38 can effectively elicit signaling through complexes that contain either the type I receptor Sax or the type II receptor Wit, and exhibits high levels of signaling in the presence of both. Furthermore, when NS cleavage is blocked in the presence of Sax and Wit, the uncleaved prodomain increases Gbb15 signaling activity. Finally, processing at S1/S0 (Gbb15) is required for activation of receptor complexes consisting of Tkv and Punt.

*Cleavage of NS or S1/S0 is sufficient for activity of Gbb-Dpp heterodimers*—BMP family members can associate and form active heterodimers (44). Given that Gbb-Dpp heterodimers are thought to contribute to patterning and differentiation of the developing wing (22–24), and that Gbb38 is the most abundant ligand form in the larval wing imaginal disc (1), we examined whether alternative Gbb processing may impact Gbb/Dpp heterodimer formation and activity. HA-tagged *dpp* (*dpp-HA*) was co-expressed in S2 cells with different *gbb-HA* wild type and cleavage mutant constructs. The composition of Gbb and Dpp protein products secreted into the media was examined using α-HA on non-reducing Western blots (Fig. 5A). When we compared secreted products produced by cells expressing *gbb-HA* wild type or cleavage mutant constructs with *dpp-HA* versus *gbb-HA* constructs alone, we detected three new bands corresponding to heterodimers composed of various Gbb cleavage products and the processed C-terminally derived Dpp ligand domain. A Gbb15-Dpp heterodimer is present when wild type *gbb* or *gbb* ^*mNS*^ are co-expressed with *dpp*. Blocking NS cleavage (*mNS*) appears to increase the abundance of the Gbb15-Dpp heterodimer. A Gbb38-Dpp heterodimer is enriched in media from cells expressing the S1/S0 cleavage mutant (*gbb* ^*mS1mS0*^). And when all three Gbb processing sites are mutated, a proGbb-Dpp heterodimer is evident. Cleaved Dpp can be secreted as a heterodimer with Gbb15, Gbb38, or proGbb. We note that when *dpp-HA* is expressed alone, Dpp-HA is not detected in media, but is instead observed in cell lysates at low abundance (data not shown). Therefore, the relatively high abundance of Gbb-Dpp heterodimers does not necessarily indicate preferential heterodimer formation or production, and instead may reflect increased secretion or decreased turnover.

**Figure 5:**
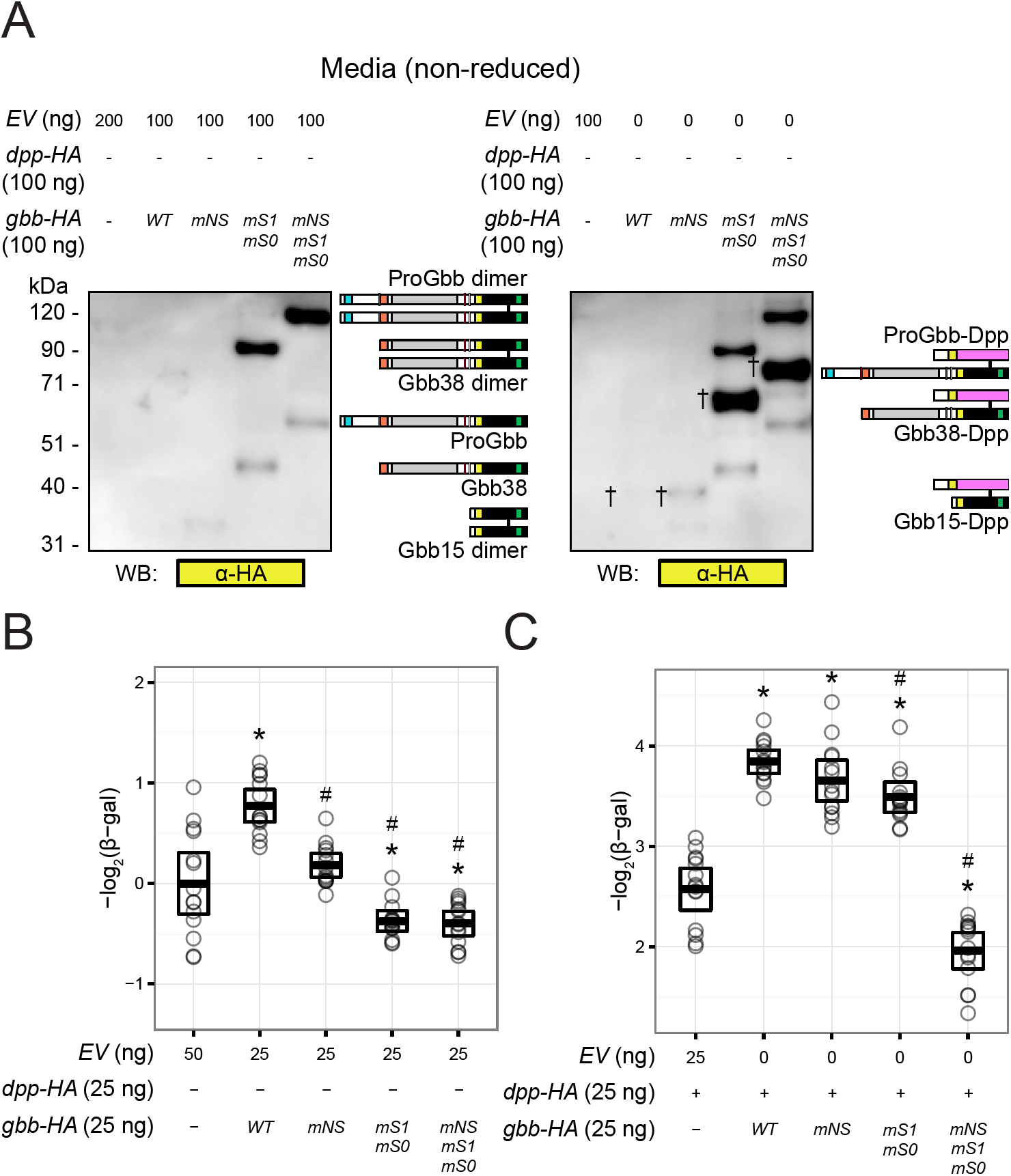
Dpp signaling activity is enhanced by forming heterodimers with Gbb cleavage mutants. A, non-reducing Western blots of media from cells co-expressing *dpp-HA* and *gbb-HA*, showing secreted homodimers, and heterodimers (indicated with). B, C, steady-state BrkSE-LacZ signaling assay, measuring activity of *gbb-HA* cleavage mutants co-expressed with *EV* (B) or *dpp-HA* (C). All signaling experiments were performed in parallel, and are presented separately to emphasize the relative effects of gbb-HA expression. Bars indicate mean and 95% CI. * indicates p < 0.05 compared to EV, # indicates p < 0.05 compared to WT *gbb-HA*, using GLHT multiple comparison test.

Next, we measured the activity of the ligands secreted from cells expressing each *gbb* construct co-expressed with *dpp* or empty vector, using the *BrkSE-LacZ* signaling assay (Fig. 5B). Similar to our previous observations, blocking NS cleavage (*mNS*) reduces signaling activity of Gbb, and blocking S1/S0 cleavage (*mS1mS0*) or all three cleavage sites (*mNSmS1mS0*) reduces activity below endogenous levels (Fig. 1E, 5B). *dpp* alone produces significantly higher signaling activity than *gbb* alone (Fig. 5B, C), as also seen when each is overexpressed in the developing wing (20), and when co-expressed a higher level of signaling activity is achieved. Whereas expression of *gbb* ^*mNS*^ alone fails to induce signaling, when co-expressed with *dpp*, signaling activity is equivalent to the levels produced by coexpression of *dpp* and wild-type *gbb*. Based on the cleavage products produced by *gbb* ^*mNS*^ (Gbb15 + NH_3_-S1; Fig. 1, S2) and the ligands detected in media from the co-expression of *gbb* ^*mNS*^ and *dpp* (Gbb15-Dpp; Fig. 5A), we attribute the higher level of signaling to Gbb15-Dpp heterodimers. Since the uncleaved Gbb prodomain (NH_3_-S1) is present in the media of cells co-expressing *gbb* ^*mNS*^ and *dpp* (data not shown), we also conclude that the Gbb prodomain does not compromise Gbb15-Dpp signaling. Whereas the production of Gbb38 failed to induce signaling in S2 cells and appeared to antagonize endogenous BMP signaling following the expression of *gbb* ^*mS1mS0*^ (Fig. 5B), we found that the co-expression of *gbb* ^*mS1mS0*^ and *dpp* produced a high level of signaling activity, likely due to the formation of Gbb38-Dpp heterodimers (Fig. 5C). Finally, when all Gbb cleavage sites are mutated (*gbb* ^*mNSmS1mS0*^) the ability of *dpp* to induce signaling is inhibited, most likely as a result of sequestering Dpp into inactive proGbb-Dpp heterodimers. Thus, cleavage of Gbb at S1/S0 or at NS enables the formation of heterodimers with Dpp leading to significant signaling activity.

*Gbb NS cleavage is required for wing vein patterning*—Mutations in the Gbb NS cleavage site have different effects on signaling activity in S2 cells, depending on whether *gbb* is expressed alone or coexpressed with *dpp*. Our lab previously observed that mutations in the Gbb NS cleavage site led to wing vein defects, including ectopic posterior cross vein (PCV) spurs (1). To generate animals that lack cleavage at the NS site we made use of a mutant form of a *gbb* genomic rescue construct (*gbbR* ^*mNS*^) in a *gbb* ^*1*^ null mutant background. Development of the PCV requries both *gbb* and *dpp*, likely acting as Gbb-Dpp heterodimers (24, 25). We assessed the requirement for the NS cleavage in Gbb-Dpp heterodimer function by making use of a sensitized genetic background where overall BMP gene dosage was reduced while preserving ligand stoichiometry. A single functional copy of *gbb* was provided by *gbb* ^*R*^ in a homozygous *gbb* ^*1*^ null in the context of reduced *dpp* function in animals heterozygous for *dpp* ^*d12*^ or *dpphr4* mutations 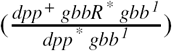. *dpp* ^*d12*^ disrupts the 3’ cis-regulatory sequences required for *dpp* expression in the wing imaginal disc (45, 46) and *dpp* ^*hr4*^ exhibits reduced protein function (47).The effect of blocking NS cleavage was determined by comparing *gbbR* and *gbbR* ^*mNS*^ in each *dpp* mutant background.

Wings of adult 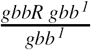 exhibit normal patterning, while 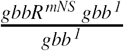 show a low penetrance of PCV phenotypes (Fig. 6A, C). Reducing *dpp* function in the context of the wild type *gbb* rescue (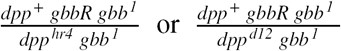) had no effect on wing patterning. In contrast, when Gbb NS cleavage is blocked in combination with *dpp* ^*hr4*^ (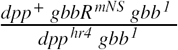) or *dpp* ^*d12*^ 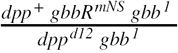 wing patterning defects, including ectopic vein formation at the tip of the 2nd longitudinal vein (L2) were significantly enhanced (Fig.6A, B). The PCV spur phenotype was enhanced in 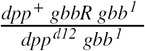(Fig. 6A, C). In summary, wing patterning defects caused by blocking NS cleavage can be enhanced by *dpp* loss of function mutations. Because Gbb NS cleavage and Gbb-Dpp heterodimers are required for PCV morphogenesis (1, 24, 25), we conclude that *in vivo* processing at the Gbb NS site is necessary for the activity of heterodimers in wing vein formation.

**Figure 6:**
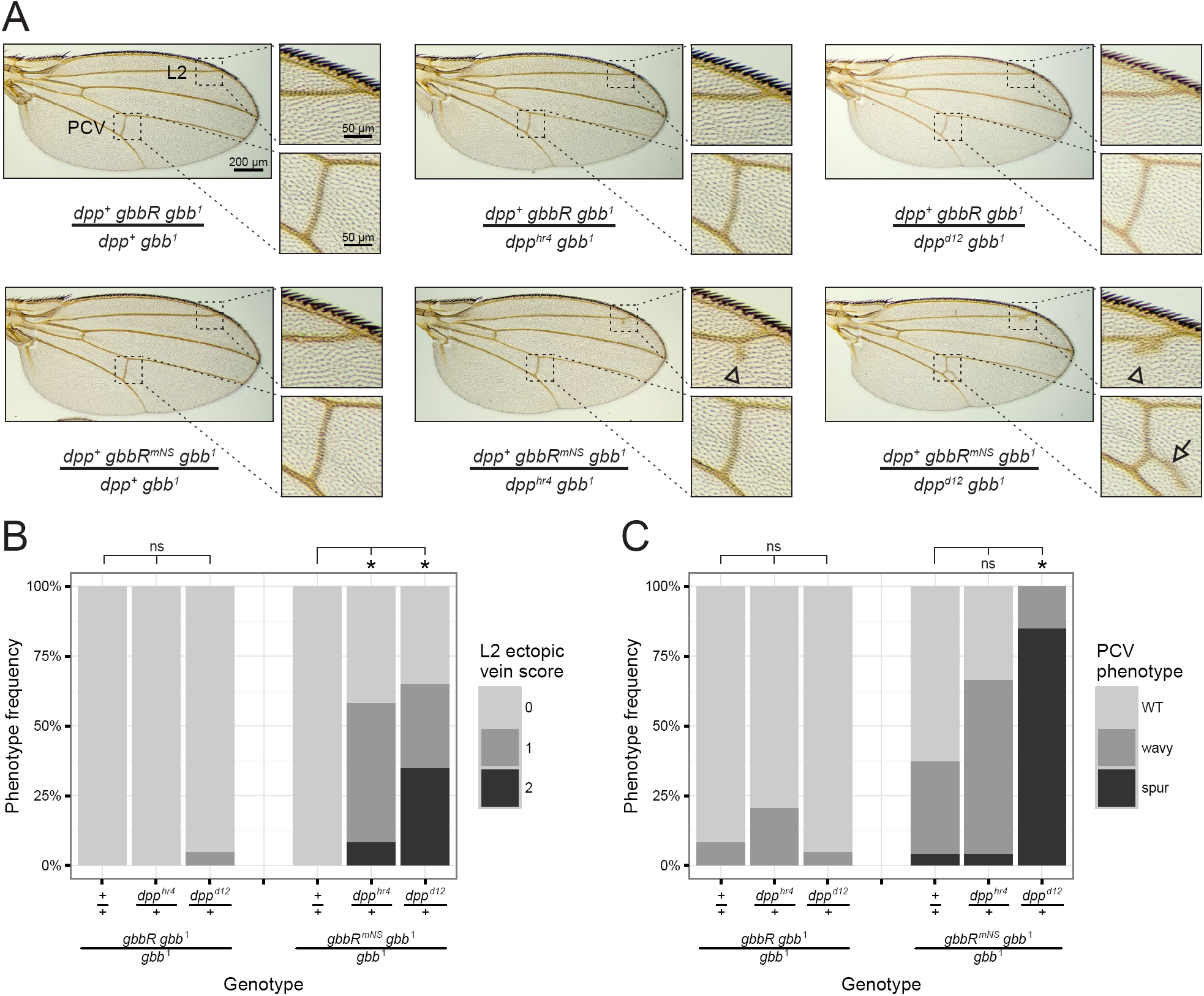
*dpp* wing phenotypes are enhanced by mutation of the *gbb* NS cleavage site. A, representative images of wings showing phenotypes resulting from genetic interactions between *gbbR*^*mNS*^ and *dpp*^*hr4*^ or *dpp*^*d12*^. Inset images show the distal tip of L2 with ectopic veins (arrowhead), and the PCV with spurs (arrow). B, proportion of L2 ectopic veins. Phenotype is scored 0 for WT, 1 for a single ectopic vein spot, and 2 for multiple ectopic vein spots. C, proportion of PCV phenotypes. * indicates *p* < 0.05, ns indicates *p* >= 0.05 using Fisher’s exact test and FDR multiple comparison adjustment.

*Gbb NS cleavage is required for pupal viability*—We examined the course of *gbbR* ^*mNS*^ *gbb* ^*1*^ development from egg to adult to identify other developmental processes that may require NS cleavage. We observed significant pupal lethality which was not previously described as a *gbb* phenotype. We scored the pupal viability of a *gbb* allelic series and found that *inter se* crosses between mild loss of function alleles (*gbb* ^*^3^*^, *gbb* ^*^4^*^, or *gbb* ^*^5^I*^) or with the *gbb* ^*1*^ null allele result in different amounts of pupal lethality (Table 1). Notably, *gbbR* ^*mNS*^ *gbb* ^*1*^ exhibits higher pupal lethality than 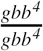 and 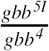. However, 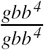 and 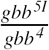 *5I* have more severe wing patterning defects than *gbbR* ^*mNS*^ *gbb* ^*1*^, including loss of the PCV and distal portions of the L4 and L5 longitudinal veins (1, 21, 23). Therefore, we hypothesized that the functions of *gbb* during pupal development require NS cleavage. Most of the dead *gbbR* ^*mNS*^ *gbb* ^*1*^ pupae exhibited developmental arrest at early stages of pupation, prior to head eversion and pigment development (48) (Fig. 7A). *gbbR* ^*mNS*^ *gbb* ^*1*^ pharates, which have undergone metamorphosis, failed to fully extend their legs (Fig. 7A, B). The failure in both head eversion and leg extension is indicative of a defect in pupal ecdysis behaviors during pupation, the process by which anatomical structures are inflated to take on their adult form (49–51). Additionally, some *gbbR* ^*mNS*^ *gbb* ^*1*^ pharates arrested at late developmental stages, just prior to or during the process of exiting the pupal case. Thus, processing of proGbb at the NS site is required *in vivo* for functions of *gbb* during pupal ecdysis and development.

**Table 1:** Pupal lethality of *gbb* mutant allelic series

| Genotype | Pupal lethality (%) | *n* |
| --- | --- | --- |
| *gbb*<sup>51</sup>/*gbb*<sup>4</sup> | 27 | 453 |
| *gbb*<sup>4</sup>/*gbb*<sup>4</sup> | 57 | 361 |
| *gbb*<sup>51</sup>/*gbb*<sup>3</sup> | 83 | 273 |
| *gbb*<sup>4</sup>/*gbb*<sup>3</sup> | 88 | 278 |
| *gbb*<sup>4</sup>/*gbb*<sup>1</sup> | 99 | 212 |
| *gbb*<sup>3</sup>/*gbb*<sup>3</sup> | 100 | 148 |
| *gbb*<sup>51</sup>/*gbb*<sup>1</sup> | 100 | 183 |
| *gbbR gbb*<sup>1</sup> | 1.4 | 349 |
| *gbbR*<sup>mNS</sup> *gbb*<sup>1</sup> | 69 | 368 |

**Figure 7:**
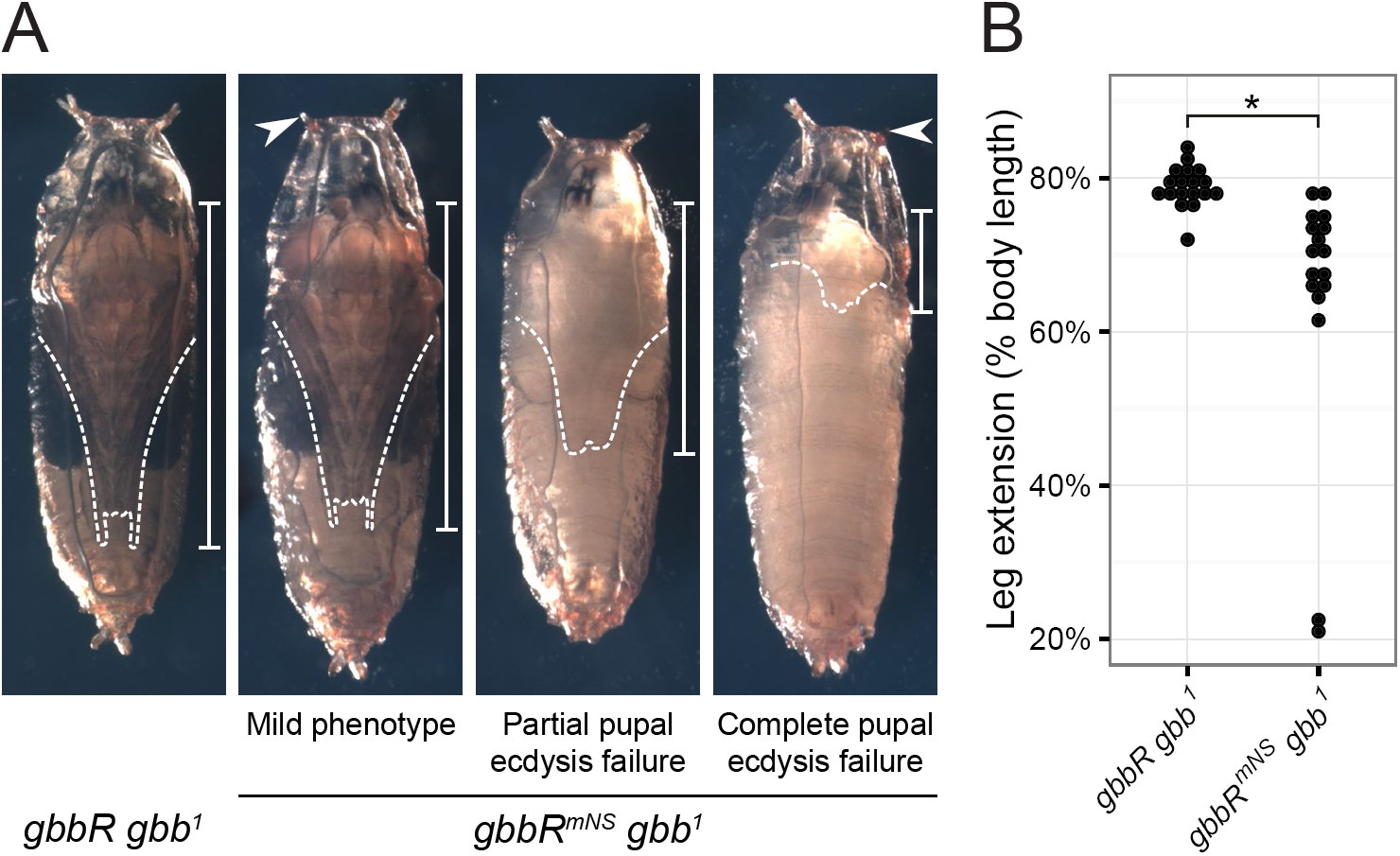
Blocking NS cleavage causes pupal ecdysis defects. A, representative images of pupae four days after pupariation. *gbbR*^*mNS*^ *gbb*^*1*^ pupal phenotypes include failure to evert anterior spiracles (arrowhead), reduced leg extension (dashed lines and bars), partial or complete failure of pupal ecdysis, and developmental arrest. B, quantification of pupal leg extension, measured as fraction of pharate body length. * indicates *p* < 0.001 using Wilcoxon rank-sum test.

*Gbb NS cleavage is required for expression of the pupal ecdysis hormone CCAP*—Pupal ecdysis involves a series of bodily movements that result in shedding of the larval cuticle, eversion of the head and appendages, and extension of the everted wings and legs to reach their adult shape (49– 51). This behavioral sequence is coordinated by hormones produced by neurosecretory neurons and endocrine cells (52). The neuropeptide Crustacean Cardioactive Peptide (CCAP) is expressed in a specific set of neurons in the ventral nerve cord (VNC) of the central nervous system; ablation of these neurons, or mutations in *ccap*, produces pupal ecdysis defects (51, 53). It has been shown that *wit* is required for pupal ecdysis and for the expression of CCAP and other ecdysis regulating hormones in CCAP neurons (29). Similarly, it has been shown that the presence of nuclear pMad in CCAP neurons is dependent on *wit*, and that the number of CCAP-expressing neurons is also reduced in *gbb* ^*1*^ 1st instar larvae. Since *gbbR* ^*mNS*^ *gbb* ^*1*^ pupal phenotypes resemble those of *wit* mutants or CCAP neuron ablation, we hypothesized that Gbb NS cleavage is required for *wit*-dependent CCAP expression. We assayed for the expression of CCAP in *gbbR gbb* ^*1*^ or *gbbR* ^*mNS*^ *gbb* ^*1*^ wandering 3rd instar larvae using *CCAP-Gal4, UAS-GFP* (*CCAP>GFP*), and used nuclear pMad immunostaining to measure BMP signaling activity in CCAP neurons.

CCAP neurons are comprised of several distinct classes of neurons in late 3rd instar larvae just prior to pupariation: interneurons (CCAP-IN), efferent neurons (CCAP-EN), late efferent neurons (CCAP-ENL), and posterior lateral neurons (CCAPPL) (Fig. 8A) (54). Since CCAP-IN neurons are not required for pupal ecdysis, and do not exhibit regulation of CCAP expression by BMP signaling, we focused our analysis on the other classes of CCAP expression neurons. Of these, only the CCAP-ENL and CCAP-PL neurons are required for pupal ecdysis. CCAP-EN and -ENL neurons are marked with nuclear pMad and Dachshund (Dac), while CCAP-PL neurons are marked with only nuclear pMad. In the ventral nerve cord (VNC) of wandering 3rd instar *gbbR gbb* ^*1*^ larvae, we detect CCAP-EN and CCAP-ENL neurons marked with nuclear pMad and Dac (pMad+ Dac+), and CCAP-PL neurons marked only with nuclear pMad (pMad+ Dac-) (Fig. 8B). CCAP expression has not been examined in *gbb* mutants at stages beyond 1st instar larvae (29), therefore, we first assessed the expression of *CCAP>GFP* in the VNC of *gbb* ^*1*^ null mutant 3rd instar larvae. CCAP-PL and CCAP-ENL neurons were almost completely undetectable in *gbb* ^*1*^ 3rd instar larvae (Fig. S5A, B). Wild-type numbers of CCAP-EN neurons could be identified in *gbb* ^*1*^ VNCs, but the expression of *CCAP>GFP* and nuclear pMad in the CCAP-EN neurons was greatly reduced (Fig. S5C, D). Therefore, we conclude that Gbb signaling is required for Wit-dependent expression of CCAP in CCAP-EN neurons. This indicates that either *gbb* is required for CCAP expression in these neurons or the developmental delay associated with *gbb* ^*1*^ nulls (21) prevented the mutant larvae from progressing to the late 3rd instar when CCAP expression begins in the CCAP-PL and CCAP-ENL neurons (54).

**Figure 8:**
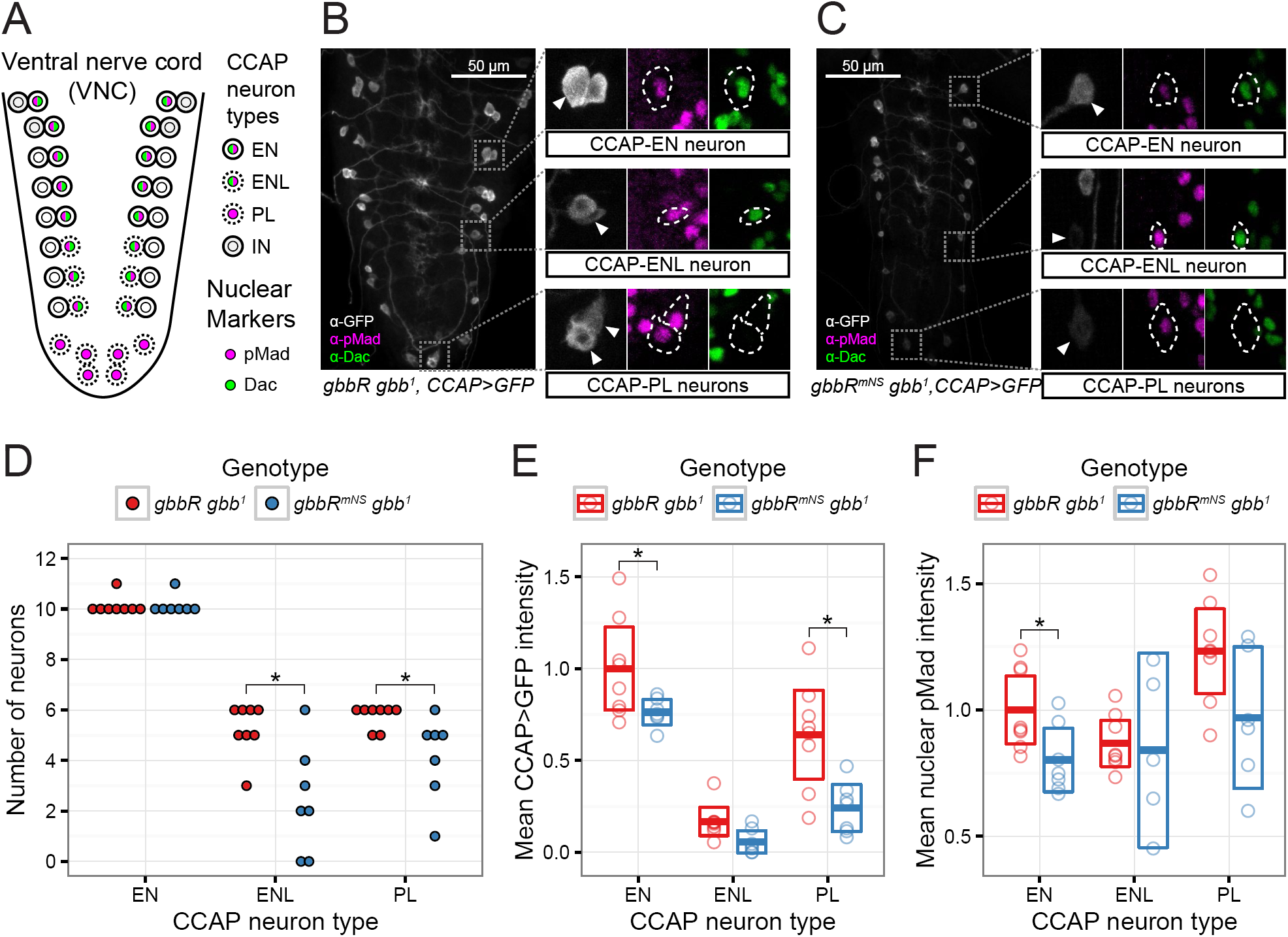
Gbb NS cleavage is required for expression of the ecdysis-regulating hormone CCAP. A, schematic of CCAP-expressing peptidergic neurons in the larval VNC, adapted from Veverytsa and Allan (54). Prior to wandering stage 3rd larval instar wandering stage, CCAP is expressed in a subset of pMad-, Dac-interneurons (IN), and pMad+, Dac+ efferent neurons (EN). At later stages, CCAP expression is activated in additional pMad+, Dac+ efferent neurons (ENL) and pMad+ posterior lateral neurons (PL). B, C, representative confocal image stacks of VNC from wandering 3rd instar larvae. CCAP>GFP is used as a reporter of CCAP expression, and nuclear Dac and pMad serve as markers and indicators of BMP signaling. Insets, CCAP neurons with pMad+ nuclei are indicated with arrowheads and dashed outline. D, number of late CCAP neurons per VNC. * indicates *p* < 0.05 using Wilcoxon rank-sum test and FDR multiple comparison adjustment. E, quantification of mean cell body CCAP>GFP signal intensity for each VNC and neuron class. F, quantification of mean nuclear pMad signal intensity for each VNC and neuron class. E, F, bars indicate mean and 95% CI, * indicates *p* < 0.05 using GLHT multiple comparison test. D-F: n = 8 VNCs for *gbbR gbb*^*1*^, n = 7 VNCs for *gbbR*^*mNS*^ *gbb*^*1*^.

To determine the consequences of blocking NS cleavage during pupal development, we examined the expression of CCAP in the VNC of *gbbR* ^*mNS*^ *gbb* ^*1*^ late 3rd instar larvae. We detected CCAP-EN neurons, and the level of *CCAP>GFP* expression in the CCAP-EN neurons was significantly reduced (Fig. 8C, E). In *gbbR* ^*mNS*^ *gbb* ^*1*^ VNCs, we found significantly fewer CCAP-PL and CCAP-ENL neurons with detectable *CCAP>GFP* expression, compared to *gbbR gbb* ^*1*^ (Fig. 8B-D). Of the remaining, detectable CCAP neurons, there was a significant reduction of *CCAP>GFP* expression in CCAP-PL neurons (Fig. 8E). We quantified the signal intensity of nuclear pMad in CCAP neurons in *gbbR* ^*mNS*^ *gbb* ^*1*^ VNCs, and compared to *gbbR gbb* ^*1*^, we observed a reduction in BMP signaling in CCAP-EN neurons (Fig. 8F). In detectable CCAP-ENL and CCAP-PL neurons with visible *CCAP>GFP* expression, we did not see a significant change in nuclear pMad intensity. In summary, we find that NS cleavage of Gbb is required to activate pMad in a subset of CCAP neurons and for the *wit*-dependent expression of CCAP. Since Gbb38 can activate Wit in S2 cells (Fig. 4A), and *gbbR* ^*mNS*^ *gbb* ^*1*^ shows defects in *wit*-dependent signaling, we propose that NS cleavage is required *in vivo* for production of Gbb38 as a ligand that preferentially activates Wit.

*Cleavage of the Gbb S1/S0 site is blocked by O-linked glycosylation*—The endogenous abundance of Gbb38 and Gbb15 varies between tissues, which suggests that that the production of Gbb38 could be a regulated process whereby S1/S0 cleavage is prohibited and only NS cleavage is permitted. O-glycosylation has been shown to prevent PC processing (55). Therefore, we considered the possibility that the modification of residues near Gbb PC sites could influence the production of one ligand form over the other. Based on NetO-Glyc 4.0 (56), proGbb is predicted to contain O-glycoslyation sites near its PC cleavage sites (Fig. 9A). Of particular note is the cluster of six high-score predicted glycosylation sites within 3 residues of the S1/S0 sites. Gbb38 is the most abundant Gbb ligand form in most 3rd instar larval tissues (1). Therefore, we used an *in vitro* cleavage assay to determine whether proGbb or Gbb38 isolated from 3rd instar larvae could be cleaved at the S1/S0 site. An HA-tagged *gbb* rescue transgene was inserted in a *gbb* ^*1*^ null background, generating *gbbR* ^*HA*^ *gbb* ^*1*^ and ensuring that Gbb-HA would be produced at endogenous levels. Gbb-HA was immunoprecipitated from lysates of *gbbR* ^*HA*^ *gbb* ^*1*^ wandering 3rd instar larvae. Similar to our previous observations, proGbb was more abundant than Gbb38, and Gbb15 was not detectable on Western blots (Fig. 9B). Treatment with recombinant Furin resulted in a decrease of proGbb and an increase of Gbb38 indicating an NS cleavage. No Gbb15 resulting from an S1/S0 cleavage was detected. To determine whether S1/S0 cleavage may be blocked by O-glycosylation, immunoprecipitated Gbb was treated with glycosidases prior to the addition of Furin. In *Drosophila*, Core 1 glycans represent the most abundant form of O-glycans, while most of the remaining O-glycans consist of a HexNAc or GlcA extension of the Core 1 glycan (57). The unmodified Core 1 O-glycan can be removed with O-glycosidase, and most extensions of the Core 1 O-glycan can be removed with exoglycosidases β-N-acetylhexosaminidase (β-HexNAcase) and β-glucuronidase (β-Glcase). Treatment of Gbb with O-glycosidase alone had no effect on Furin cleavage. However, following treatment with all three glycosidases, addition of Furin produced Gbb15. We conclude that in *Drosophila* 3rd larval instar, S1/S0 cleavage of Gbb is blocked by an extended O-glycan, resulting in the production of Gbb38 by cleavage of the NS site. Since S1/S0 cleavage occurs in S2 cells, it is conceivable that specific glycosyl-transferases are expressed in 3rd instar larvae that are responsible for the O-glycosylation of the Gbb S1/S0 site, and that these enzymes are not expressed in S2 cells. Thus, S1/S0 cleavage is allowed and Gbb38 is not produced as a mature ligand by S2 cells.

**Figure 9:**
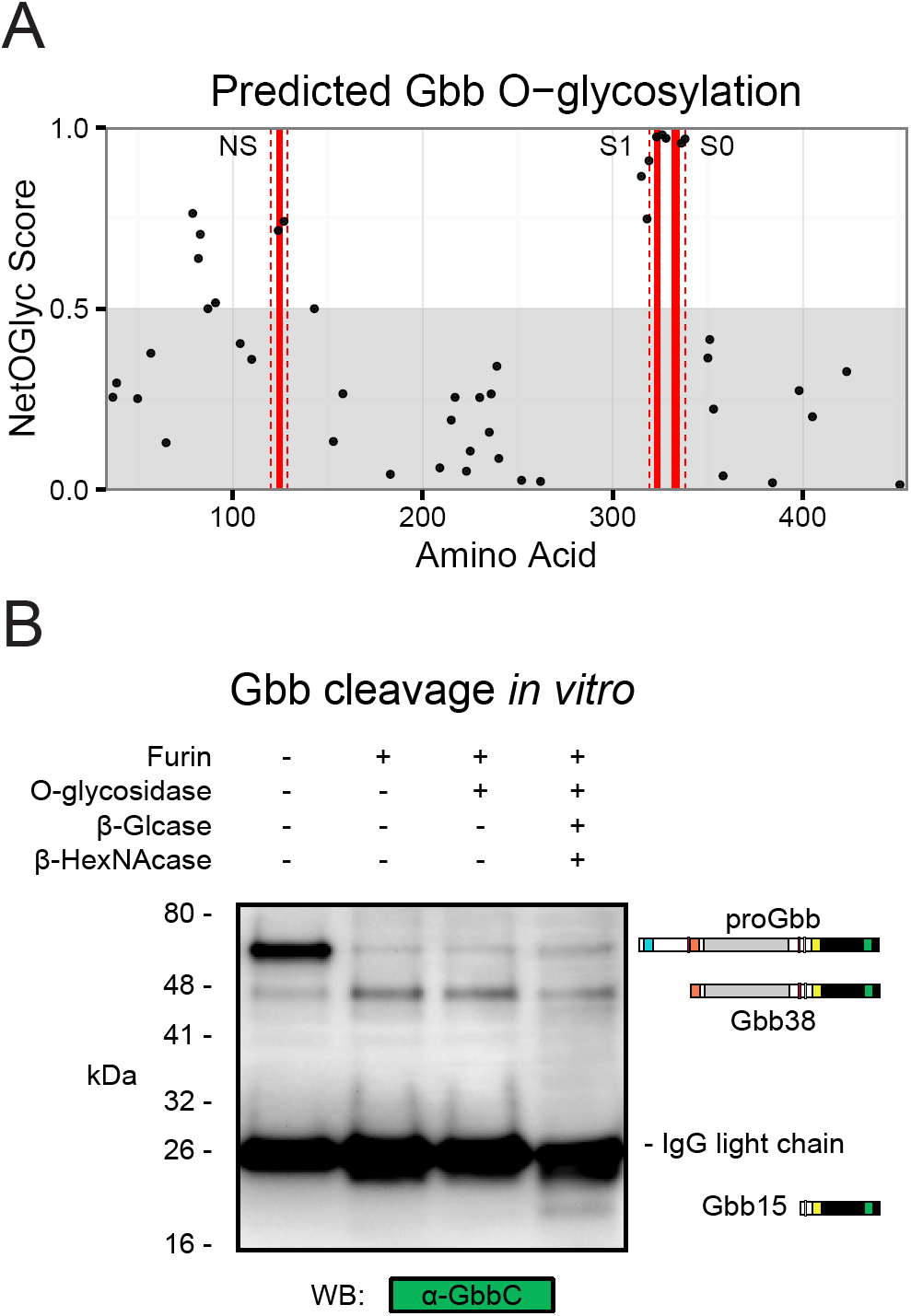
O-linked glycosylation blocks S1/S0 cleavage and Gbb15 production. A, predicted O-linked glycosylation sites on Ser/Thr residues in Gbb. Residues with NetOGlyc 4.0 scores >= 0.5 indicate predicted O-linked glycosylation, and scores < 0.5 (grey) indicate no predicted glycosylation. PC cleavage sites are indicated with solid red lines, and dashed red lines indicate ±3 residues. B, *in vitro* cleavage of Gbb-HA immunoprecipitated from *gbbR*^*HA*^ *gbb*^*1*^ 3rd instar larvae. Immunoprecipitate was treated with recombinant Furin, with or without O-glycosidase, β-glucuronidase (β-Glcase), or β-N-acetylhexosaminidase (β-HexNAcase), and analyzed on a non-reducing Western blot.

## DISCUSSION

### Regulation of Gbb signaling by alternative cleavage

Alternative processing of the Gbb proprotein can produce ligands of different sizes, Gbb38 and Gbb15 (Fig. 10). These ligands vary by the inclusion of the Core region of the prodomain, which appears to confer receptor preference. In S2 cells, Gbb15, generated by PC processing at the S1/S0 site, is secreted as a loosely associated complex with the two prodomain fragments (NH_3_-NS and NS-S1) produced by cleavage at the NS site. In this configuration, Gbb15 can activate signaling via either type II receptor, Punt or Wit. When NS cleavage is blocked, Gbb15 forms a tightly associated latent complex with the uncleaved prodomain (NH_3_-S1) with little Punt-mediated signaling activity. In this case, more of the resulting Gbb15 is found in the media than in wild type when NS cleavage occurs, suggesting that the Gbb15-uncleaved prodomain latent complex influences either protein turnover, secretion, and/or binding to cell surface receptors that can also affect ligand turnover. Interestingly, when S2 cells are induced to express Wit, the Gbb15-uncleaved prodomain latent complex is not restricted in its ability to signal and exhibits full activity. In the absence of S1/S0 cleavage, NS-cleaved Gbb38 is secreted in complex with the small N-terminal fragment of the prodomain (NH_3_-NS) and is only able to signal through Wit. These findings indicate that the Gbb prodomain influences type II receptor usage. Signaling through Punt is restricted by either the Core domain as part of Gbb38, or by the intact prodomain when it is non-covalently associated with Gbb15, while Wit can mediate all ligand-prodomain cleavage fragment combinations.

**Figure 10:**
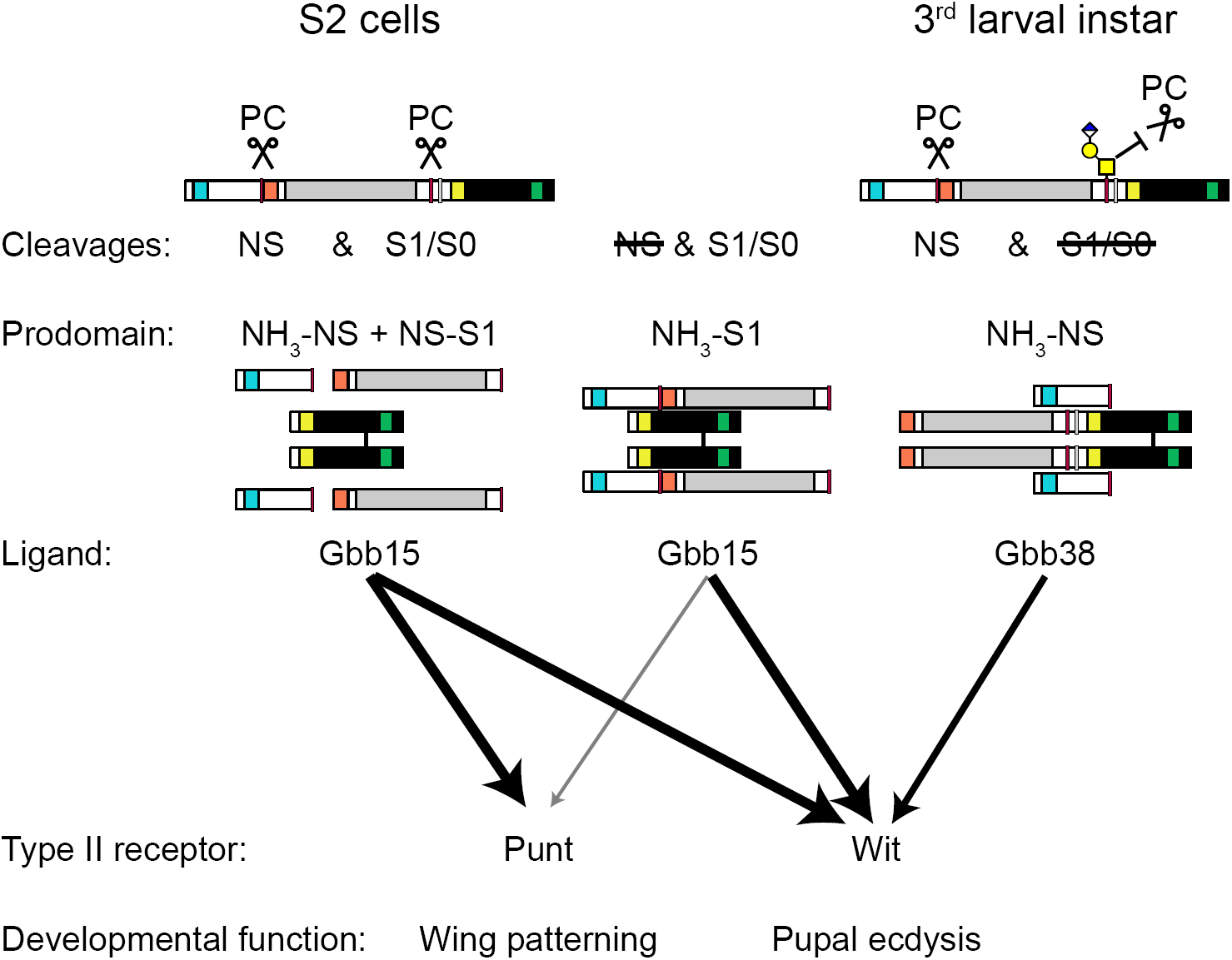
Model of Gbb processing, prodomain-ligand complexes, and activity. Gbb expressed in S2 cells is cleaved by a proprotein convertase (PC) at both the NS and S1/S0 site, and the resulting ligand (Gbb15) and prodomain cleavage products (NH_3_-NS and NS-S1) are secreted as a loosely associated complex that has signaling activity mediated by Wit or Punt. When NS cleavage is blocked by mutation of the NS site (mNS), the uncleaved prodomain (NH_3_-S1) is tightly associated with Gbb15, and Punt-mediated signaling is reduced. When S1/S0 cleavage is blocked, by mutation of the S1/S0 site (mS1mS0), the ligand Gbb38 is secreted in complex with the N-terminal prodomain cleavage fragment (NH_3_-NS) and Punt-mediated signaling is greatly reduced. In the 3rd larval instar, cleavage of the S1/S0 site is blocked by O-glycosylation, depicted here as hypothetical extended Core 1 O-glycan. Thus, Gbb15 production is blocked, and Gbb38 is produced as a ligand that is required for Wit-mediated regulation of pupal ecdysis.

The abundance of Gbb38 and Gbb15 varies in different tissues of the developing animal, suggesting that either the production or turnover of the two ligand types is regulated. Here, we show in 3rd instar larvae that proteolytic processing at the S1/S0 can be influenced by O-glycosylation leading to preferential processing at the NS and an abundance of Gbb38 (Fig. 10). Furthermore, we find that NS cleavage is required *in vivo* for wit-dependent functions in pupal ecdysis and the regulation of CCAP hormone in specific CNS neurons that are critical for metamorphosis. We propose that at least one mechanism responsible for regulating the alternative cleavage of Gbb involves O-glycosylation of residues near the S1/S0 site, resulting in the production of Gbb38 as a ligand that preferentially activates Wit.

### Gbb NS cleavage is necessary for Gb-b-Dpp heterodimer activity patterns

We found that 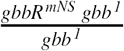 adult wings have a low occurrence of vein patterning defects. However, in combination with *dpp* loss of function mutations (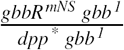) a substantial enhancement of defects is apparent, including PCV spurs and ectopic vein material. These phenotypes do not resemble those displayed by loss of function mutations in *gbb* or *dpp* which typically cause the absence of longitudinal veins and cross veins, rather than the formation of ectopic veins (21–23, 58). Instead, the wing phenotypes of 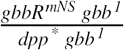 resemble those caused by mutations in or overexpression of several secreted BMP extracellular regulators. During PCV morphogenesis, Gbb/Dpp heterodimers expressed in longitudinal vein primordia are bound by the extracellular regulators Crossveinless (Cv) and Short gastrulation (Sog), the Drosophila Chordin ortholog (59). When complexed with Cv and Sog, the Gbb/Dpp heterodimer is inactive. However, Crossveinless-2 (Cv-2) serves to localize the complex to the PCV primordia between longitudinal vein L4 and L5 where cleavage by the Tolloid (Tld) metalloprotease releases Gbb/Dpp for signaling. Together, these extracellular regulators comprise a system that shuttles Gbb/Dpp heterodimers from the longitudinal veins to the PCV primordia. Experimental disruption of the stoichiometry or spatial distribution of these extracellular regulators produces PCV spurs and ectopic veins (26, 60–62) similar to those that we see with the loss of Gbb NS cleavage in the context of a reduction of *dpp* dosage. Similarly, loss of function mutations of another regulator of BMP signaling, *larval translucida* (*ltl*), produces the same phenotype (63). Structural studies of TGF-β family proteins complexed with prodomains or extracellular regulators suggest that their binding to BMPs could be mutually exclusive (6). Since blocking NS cleavage increases prodomain association with ligands, we suggest that the uncleaved prodomain could bind Gbb-Dpp heterodimers and interfere with the functions of Sog, Cv, or other extracellular regulators.

### Functions of Gbb in regulation of ecdysis hormones

We demonstrate that NS-cleaved Gbb is required during metamorphosis for the proper expression of hormones that control pupal ecdysis, the process by which imaginal tissues assume their adult form. In addition to the pupal ecdysis defects we described, we have also observed failure of eclosion, wing inflation, and defects in tanning whereby newly emerged adults remain soft for many hours after heterozygous siblings have hardened their cuticle (data not shown). We also note that the *gbbR* ^*mNS*^ *gbb* ^*1*^ pupal phenotype includes failure to develop pigment, but pigment development is not affected by ablation of CCAP neurons or mutations in CCAP and related hormones (51, 53). Our laboratory has also observed larval ecdysis defects in *gbb* loss of function mutants, including significant developmental delays and failure to shed cuticle during each molt^1^. Together, these phenotypes all suggest that *gbb* has a broader and previously unrecognized role in the regulation of ecdysis in both larval and pupal development. Regulation of ecdysis by *gbb* could be related to its role in the regulation of energy homeostasis (64), which is intimately tied to the organism’s ability to undergo or delay ecdysis (52), or to the role of *gbb* in circadian neural circuit development (65, 66). With regard to the regulation of pupal ecdysis hormones, *wit* signaling is required specifically in CCAP neurons for CCAP expression (29). 1st larval instar *gbb* ^*1*^ have fewer CCAP expressing neurons, and a resulting reduction in the expression of other ecdysis hormones, Myoinhibiting Peptide and Bursicon β (29). Since NS cleavage of Gbb is required for the regulation of CCAP, it is conceivable that NS cleavage is required for the production of active Gbb38 when S1/S0 cleavage is blocked. Gbb38 may be produced as a ligand that preferentially activates Wit in the nervous system, and acts to regulate hormonal expression throughout development.

### Regulation of receptor binding and preference by Gbb and TGF-βfamily prodomains

Our data suggests that the Core domain of the Gbb prodomain influences the preferential use of type II receptors. When the Core domain fragment (NS-S1) is non-covalently associated with Gbb15, signaling activity is permitted with any receptor. In contrast, when the Core domain remains attached to the C-terminal ligand domain (in Gbb38) or to the NH_3_-terminal domain (in the NH_3_-S1 uncleaved prodomain), signaling via the type II receptor Punt is inhibited, while signaling via the type II receptor Wit is permitted. These findings are consistent with previous reports that have found that the prodomain from several TGF-β family members can interact with type II receptor binding surfaces on the ligand. The Core/Arm domain occupies the type II receptor binding surfaces of the mature ligand in the crystal structures of ligand-prodomain complexes of TGFβ1, Activin A, and BMP9 (10, 18, 67). The type II receptors ACVR2A/B/Punt and BMPR2/Wit primarily differ by the sequence and conformation of the "A loop", which is adjacent to receptor- and prodomain-binding sites on the mature ligand, and may change conformation upon ligand binding (6, 68). We speculate that the A loop determines the relative ability of each type II receptor to displace prodomains from ligands.

Competition for ligand-binding between type II receptors and prodomains has been demonstrated *in vitro* for BMP7, BMP9, Activin A, Inhibin A, and AMH (17–19, 69–71). However, until now, the *in vivo* consequences of prodomain-receptor competition have not been established. While type II receptor usage depends on the alternative cleavage of Gbb, the prodomains of other BMPs and TGF-β family ligands may be able regulate type II receptor preference *in vivo* even without alternative cleavage. For example, the differential expression of type II receptors with varying abilities to displace prodomains might explain why the BMP10 ligand-prodomain complex is latent in C2C12 myoblasts, but not in multiple endothelial cell lines (12, 72).

We found that when NS cleavage is blocked, Gbb forms a latent prodomain-ligand complex that has reduced Punt-dependent signaling activity. Though we observed this latency by mutating the NS cleavage site, cleavage at this site could be regulated *in vivo* by an as yet unknown mechanism. Latent Gbb could be produced when NS cleavage is blocked by predicted O-glycosylation. Latency of Gbb (Gbb15 + uncleaved prodomain) represents a novel finding for the BMP5/6/7/8 sub-family, which have been shown to form active prodomain/ligand complexes (12, 17, 73). Instead, the activation of Gbb by NS cleavage is analogous to the activation of latent GDF8, GDF11, and BMP10 complexes by Tolloid family metalloproteinases, which cleaves at a site that aligns near the NS site on Gbb (12, 39, 40). From this perspective, the functional similarity of Gbb NS cleavage and GDF8/11 Tld cleavage could represent convergent evolution of a ligand activation mechanism. Additionally, there may be other mechanisms that regulate prodomain-ligand association and latency. We saw that the Gbb prodomain-ligand association was higher at pH 6.5 than at pH 7.4, even when the prodomain is cleaved at the NS site (Fig. 3, S3, S4). This observation is intriguing because latent TGFβ1 can be activated by physiologically relevant shifts in pH (74). Although the hemolymph of *Drosophila* larvae is acidic (41), there could certainly be neutral extracellular microenvironments within developing tissues. In such an environment, neutral pH could promote dissociation of the prodomain from Gbb15 or Gbb38, to allow other protein interactions and alter signaling output.

We note that the latency conferred by the uncleaved Gbb prodomain is not absolute. In the presence of endogenously expressed receptors in S2 cells, the uncleaved prodomain reduces but does not completely abolish Gbb15 activity. However, this inhibitory effect is increased when Punt is overexpressed, and alleviated when either Tkv or Sax is overexpressed with Punt. While the Core/Arm domain of TGF-β family prodomains occupies the type II receptor binding sites, the N-terminal portion of prodomains forms an alpha helix that occupies type I receptor binding sites (10, 67, 75). Therefore, it is possible that type I receptors cooperate with Punt to effectively displace the uncleaved prodomain from Gbb15. Alternatively, NS cleavage could convert the Gbb prodomain-ligand complex from a closed, inactive form into an open, active form (18, 76).

### Regulation of TGF-β PC cleavage by O-linked glycosylation

O-glycosylation regulates cleavage of several known PC substrates (77). To our knowledge, Gbb represents the first known TGF-β family member where alternative processing is influenced by O-glycoslyation. However, there are several indications that O-glycosylation could be a more general regulatory mechanism for TGF-β family member processing. *In vitro* cleavage of peptides containing the cleavage sites of BMP7 and Inhibin α can be blocked, at least partially, by O-glycosylation (55). Like Gbb, the Inhibin α proprotein contains multiple cleavage sites. Cleavage at only the N-terminal site produces Inhibin α Nα C, a Gbb38-like protein that is produced when Inhibin is expressed in 293T cells (78). Alternative processing of Inhibin α could potentially be controlled by expression of specific O-GlcNAc transferases, which differ in their ability to block cleavage of an Inhibin α peptide *in vitro* (55). Protein processing can also by regulated by phosphorylation that blocks O-glycosylation, but not PC cleavage. The Golgi kinase Fam20C permits FGF23 cleavage by Furin, by blocking inhibitory O-glycosylation (79). Intriguingly, the PCV spur and ectopic L1 vein phenotype we observe in *gbbR* ^*mNS*^ *gbb* ^*1*^ has also been observed in mutants of the Drosophila Golgi kinase Fj (80, 81). Conceivably, Fj might be necessary to block O-glycosylation that is predicted at the NS site (Fig. 9A), and thus permit PC cleavage. Further studies are warranted to examine the potential impact of Fj and protein phosphorylation on Gbb processing and activity during pupal wing vein formation.

### Conclusion

We find that the maturation of different BMP ligand forms by alternative processing can lead to differential activation of receptors and specific signaling outputs. Cleavage of proGbb at its S1/S0 site is blocked by O-glycosylation in 3rd instar larvae, resulting in the production of Gbb38 that has preference for Wit-mediated signaling. Cleavage at the NS site is required for Gbb15 homodimer signaling via the type II receptor Punt, but not for Gbb-Dpp heterodimers. Since many other TGF-β family prodomains have predicted PC cleavage sites within their prodomains, and mutations in these predicted sites are associated with human developmental abnormalities (1), we suggest that alternative processing is an important mechanism for regulating TGF-β family signaling activity and receptor preference.

## EXPERIMENTAL PROCEDURES

### Drosophila strains and cell lines

Wild type strains used in experiments were *w* ^*1118*^ or b pr cn bw. Rescue strains *gbbR gbb* ^*1*^ and *gbbR* ^*mNS*^ *gbb* ^*1*^, produced by inserting a gbb genomic fragment at 53B2 using the øC31 system, were described previously (referred to as *gbbR-gbb* ^*\**^ *gbb* ^*1*^ in (1)). *gbbR* ^*HA*^ *gbb* ^*1*^ was generated in the same way, with an HA tag inserted in the *gbb* coding sequence between residues 351 and 352. 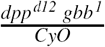and 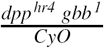 strains were created by recombination. *CCAP-Gal4*, obtained from the Bloomington Drosophila Stock Center (BDSC #25686), and *UAS-GFP* were recombined with *gbb* ^*1*^, *gbbR gbb* ^*1*^, and *gbbR* ^*mNS*^ *gbb* ^*1*^. Drosophila strains were maintained on standard cornmeal-sugar-yeast media supplemented with live yeast.

Schneider 2 (S2) cells were obtained from the *Drosophila* Genomics Resource Center (DGRC #6), and were maintained in M3 media supplemented with 10% Insect Media Supplement (Sigma I7267) and 2% FBS (Corning 35-010CV). Cells were passaged weekly, and used for up to 25 passages.

### DNA constructs

Constructs containing the *gbb* coding sequence and UTR, with *mNS* (R123G, R126G), *mS1* (R322G, R325G), and an HA tag inserted between residues 351 and 352,were described previously (1). The *mS0* mutation, K334N (2), was introduced in *pDONR221-gbb* constructs by site directed mutagenesis using primers 5’-CACGCAAGCGCAATAAGTCGGTGTCG-3’ and 5’-CGACACCGACTTATTGCGCTTGCGTG-3’. Correctly mutagenized transformants were confirmed by sequencing. Gateway cloning was used to transfer *pDONR221* (Invitrogen #12536017) inserts into *pAW* (DGRC #1127) for constitutive expression, and into *pMT-DEST48* (Invitrogen #11282018) for metallothionein-induced expression.

Expression constructs for type I receptors, *pAWF-sax* and *pAWF-tkv*, were described previously (1). Expression constructs for type II receptors, *pAWH-punt* and *pAWH-wit*, were created by in-frame insertion of *punt* and *wit* cDNA (a gift of M. O’Connor) into the *pAWH* expression vector containing a C-terminal 3xHA tag (DGRC #1096). *pAW-Dpp-HA* was constructed by site directed mutagenesis of *pAW-Dpp*, inserting a 1xHA tag between residues 485 and 486 of Dpp. Expression construct for FLAG-tagged *Mad*, *pAc-Mad-FLAG*, was a gift of M. O’Connor. *pCoPuro* was used for puromycin selection of stably transfected cells (82).

### Antibodies

Rabbit polyclonal antisera was raised against peptides corresponding to specific regions within the Gbb prodomain (Pierce Custom Antibody Services). α-GbbN was raised against peptide GKDQTIMHRVLSEDDK, corresponding to aa 46-61 of pre-proGbb. α-GbbCore was raised against peptide SADLEEDEGEQQKNFITD, corresponding to aa 127-144 of pre-proGbb. Antisera from multiple animals and batches was selected for maximum specific signal. For use in immunoprecipitations and to reduce background on Western blots, antibodies were purified by affinity to their respective peptide antigens.

Remaining primary antibodies include mouse α-Dac (Developmental Studies Hybridoma Bank (DSHB) 1-1), mouse α-FLAG (Sigma-Aldrich M2), mouse α-GbbC (DSHB 3D6-24 (1)), chicken anti-GFP (Invitrogen A10262), and rat α-HA (Roche 3F10). Rabbit α-pMad was an antibody raised against human pSmad3, which has an identical phosphorylated C-terminal SSVS motif that is not shared with the Drosophila Smad3 ortholog (Abcam EP823Y, (83)).

Secondary antibodies used were goat α-chicken-IgY:AF488 (Invitrogen A11039), goat α - mouse-IgG:AF568 (Invitrogen A11004), goat α - mouse-IgG:HRP (Jackson 115-035-146), goat α - mouse-IgGLC-HRP (Millipore AP200P), goat α - rabbit-IgG:AF633 (Invitrogen A21070), mouse α - rabbit-IgGHC:HRP (Abcam 2A9), goat anti-rat-IgG:HRP (Jackson 112-135-167). Clean-Blot (Thermo Scientific 21230) was used to specifically detect folded primary antibodies on western blots without interference from denatured antibodies used for immunoprecipitation.

### Statistical analysis

All statistical analysis was done in R 3.3.1. Data transformations were determined using the Box-Cox procedure. All statistical inference uses multiple-comparison adjusted tests. Inference of continuous numerical data was done using the general linear hypothesis test (GLHT) using the glht() function in the package multcomp (84). To control the family-wise error rate, the family of hypothesis was explicitly defined for each analysis, and included all orthogonal pair-wise comparisons including a control condition. When comparing multiple treatments with a single control this approach is identical to Dunnett’s multiple comparison test. Observations with studentized residuals > 5 were rejected as outliers. For experiments that were repeated multiple times, data was pooled and analyzed using “experiment” as a co-variate. The Wilcoxon rank-sum test was used for discrete numerical data, and Fisher’s exact test was used for categorical data, controlling for multiple comparisons using the false discovery rate (FDR) (85).

### Protein sample preparation

S2 cells were transiently transfected with *pAW-gbb-HA* or *dpp-HA* expression constructs containing the indicated cleavage site mutations, using Effectene transfection reagent (Qiagen). At 3 days post-transfection, cells were resuspended and pelleted by centrifugation. Media samples were taken from the supernatant, and 0 or 100 mM DTT and Nupage 4x LDS sample buffer (Novex) was added. Cell lysate samples were made by adding 1x LDS sample buffer with 100 mM DTT to cell pellets, to a volume equivalent to media samples. We observed significant and potentially confounding protein degradation when using Laemmli sample buffer or 95 ^°^C heating, so instead samples were heated for 65 ^°^C for 10 min using the Nupage sample buffer.

### Western blots

SDS-PAGE was run on NuPage bis-tris 12% acrylamide gels with MOPS buffer (Novex), or modified tris-tricine 10% acrylamide gels (86). Protein was transferred to PVDF membranes using a Trans-Blot semi-dry transfer apparatus (Biorad) at 10 V for one hour, with transfer buffer containing 100 mM Tris, 192 mM glycine, and 20% methanol. Membranes were blocked with 5% BSA in tris buffered saline with 0.1% Tween 20 (TBSTw), or with protein-free blocking solution (Thermo Scientific 37570) supplemented with 0.1% Tween 20. Primary antibodies and secondary antibodies were diluted in blocking buffer. Primary incubations were done shaking overnight at 4 ^°^C, followed by washes in TBSTw. Secondary incubations with HRP conjugated antibodies were done for one hour at 4 ^°^C, followed by TBSTw washes. Blots were imaged using a chemiluminescent substrate (Pierce Supersignal West Dura, #34075) and a digital imaging system (Kodak IS4000S or Biorad Chemidoc XRS). For reprobing, blots were stripped with Restore acidic glycine stripping buffer (Thermo Scientific). Bands were quantified with NIH ImageJ, using the gel tool to take lane profiles and measure the integrated peak signal.

### Immunoprecipitation

S2 cells were transiently transfected with the indicated *pAW-gbb-HA* expression construct. At 3 days post-transfection, 1 mL conditioned media was concentrated 50x with 10K MWCO centrifugal ultrafiltration units (Millipore Amicon Ultra-4), and diluted with IP buffer (25 mM Bis-Tris pH 6.5 or Tris pH 7.4, 150 mM NaCl, 1% NP40, 1 mM EDTA, 5% glycerol). Primary antibodies were added to the media concentrates and incubated 1 hr rotating at 4°C. Antibodies were pulled down with magnetic Protein G Dynabeads (Novex), washed 3x with IP buffer and 1x with water. Samples were prepared by adding 1x Nupage LDS sample buffer with 100 mM DTT, and heating for 10 minutes at 65 ^°^C. Western blots were done as previously, except using secondary detection reagents with minimal cross-reactivity to the immunoprecipitating antibodies. For α-GbbN and α-GbbCore, Clean-blot (Thermo Scientific) was used to detect natively folded primary antibodies. For α-GbbC, goat α - mouse-IgG-LC:HRP (Millipore AP200P) was used to detect specific signals without interference from the mouse IgG heavy chain, and with minimal interference from rat IgG heavy or light chains.

### Heparin affinity chromatogrpahy

Conditioned media from S2 cells expressing *gbb-HA* was loaded on a 1 mL HiTrap Heparin HP column (GE healthcare) operated with a peristaltic pump at 1.0 mL/minute. The column was washed with 10 volumes of a 10 mM sodium phosphate pH 6.0-7.4 gradient, followed by 10 volumes of a 10 mM sodium phosphate, 0-1 mM NaCl gradient. 1 mL fractions were manually collected, precipitated with TCA, and resuspended in 100 μL sample buffer for analysis by western blot.

### Furin and glycosidase assay

HA-tagged Gbb was immunoprecipitated from *gbbR* ^*HA*^ *gbb* ^*1*^ wandering 3rd instar larvae. Larvae were ground in lysis buffer (25 mM Tris pH 7.5, 150 mM NaCl, 1% NP40, 1 mM EDTA, 5% glycerol, 0.1% SDS) using a glass tissue grinder on ice. Crude lysate was centrifuged to remove cuticle, debris, and fat. Cleared lysate was incubated for 1 hour with α-HA magnetic beads (Pierce #8836). Each IP contained 100 μg α - HA beads and the lysate of 68 mg of larvae (approximately 20 larvae). Beads were washed twice with lysis buffer, and twice with Furin/Glycosidase reaction buffer (25 mM Tris pH 7.5, 50 mM KCl, 5 mM CaCl2, 0.5% Triton X-100). Samples were briefly denatured by heating for 5 minutes at 65° C with 1 mM β-ME. Samples were treated with the indicated glycosidases for 1 hour at 37 ^°^C, using 20,000 units of O-glycosidase (NEB P0733), 5 units β-N-acetylhexosamindase (NEB P0721), or 0.1 units β-glucuronidase (Sigma-Aldrich G8295). After an-other 5 min 65 ^°^C incubation, samples were treated overnight at 37 ^°^C with 4 units Furin (P8077S), and analyzed by Western blot.

### Cell signaling assays

The *BrkSE-LacZ* assay was used to measure BMP signaling activity in S2 cells (34). In summary, the LacZ reporter expression is activated by co-transfected Notch intracellular domain (NICD) and Su(H). LacZ expression is quantitatively silenced by binding of the pMad-Med-Shn complex to the *Brinker* silencer element (*BrkSE*). For signaling assays, 2 10 ^5^ S2 cells were plated in 96-well plates. Cells were transfected with 5 ng *BrkSE-LacZ*, 7 ng *NICD*, 7 ng *Su(H)*, 1 ng *pAc-luciferase* as a transfection control, and a total of 50 ng of *gbb*, *dpp*, or receptor expression constructs, using Effectene transfection reagent (Qiagen). At day 3 post-transfection, Luciferase and β-galactosidase activity was measured with the Dual-Light assay (Novex). Repeated assays of identical samples yielded consistent β-galactosidase activity (*R* ^2^ = 0.93), but luciferase activity was far less consistent (*R* ^2^ = 0.57), so we only include β-galactosidase activity in our analysis. Each *BrkSE-LacZ* signaling experiment used 3-6 technical replicates per condition, and was repeated three times in total. For analysis, data from all three replicates was pooled and fit to a model of the form–log(*BrkSE-LacZ*) = *β_Transfection_* + *β_Experiment_*, and GLHT was used for multiple-comparison adjusted inference of differences between transfections.

For measurement of BMP signaling activity by pMad levels, *pAc-Mad-FLAG* transfected cells were treated with conditioned media containing Gbb. Conditioned media was collected from S2 cells stably transfected with *pMT* empty vector or *pMT-gbb-HA* constructs, and induced with CuSO_4_ in serum-free media. Ligand concentration was measured by western blot. Cleavage mutant conditioned media was diluted to match wild-type ligand concentration, using *pMT* (empty vector) conditioned media. At the indicated time points, media was aspirated from centrifuged cell pellets, and cell lysate samples were prepared for western blots. Signaling activity was measured as the ratio of pMad to FLAG-tagged Mad band intensities.

### Microscopy

For measurements of wing phenotypes, 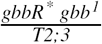*1* females were crossed with, 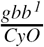, 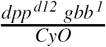 or 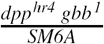 males. Wings were dissected from male and female Cy+ progeny, and mounted in Gary’s Magic Mounting Media between two cover slips, to permit imaging from both sides. The dorsal face of each wing was imaged using a Microphot FXA compound microscope (Nikon), using a Plan 4/0.13 objective (Nikon 208135) and a D600 digital SLR camera (Nikon). Images were blinded for scoring the presence and severity of wing vein abnormalities. Representative images were selected from female wings with median phenotypes and good image quality.

For measurements of pupal phenotypes, pupae were mounted on 1% agarose plates and observed each day. At 90-116 hours after pupariation, pupae were imaged from the ventral side with an oblique transmitted light source with a Ziess Lumar V12 stereo microscope and Axiocam Mrc5 camera. Images were blinded for measurements of leg extension in ImageJ, taken as the ratio between the pharate head to tail and head to leg tip length.

For examination of CCAP expression thelarval VNC, 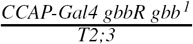was crossed to 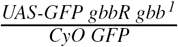, containing either the wild type or *mNS* rescue construct. Tb+, CyO GFP-wandering 3rd instar larvae were selected for dissections. Larvae were filleted and all organs that might obscure the VNC, including the lobes of the CNS, were removed. Fillets were marked according to genotype, fixed for 20 minutes in PBS plus 4% paraformaldehyde, and all genotypes were combined in a single tube. Fillets were blocked for 60 minutes with 10% NGS in PBS plus 0.3% triton x-100 (PBSTr). Primary incubation with α-GFP, α-Dac, and α-pMad was done rotating at 4 ^°^C overnight, followed by washes in PBSTr. Secondary incubation with fluorophore conjugated secondary antibodies was done rotating for one hour rotat-ing at 4 ^°^C, followed by washes with PBSTr and a 10 minute stain in 1 μg/mL Hoechst. Fillets were equilibrated overnight in 80% glycerol, 0.5% n-propyl gallate mounting media at 4 ^°^C, and mounted the next day. The VNC was imaged on a Zeiss LSM800 confocal laser scanning microscope, with a 40x/1.3 Zeiss Plan-APOCHROMAT objective. Image stacks were obtained with 0.16 μm horizontal resolution and 1.0 μm slice thickness, and blinded for analysis. For each CCAP neuron in segments T3-A9, CCAP>GFP signal intensity was quantified in cell bodies, and pMad nuclear intensity was quantified for each CCAP neuron using ImageJ. Images with approximately median numbers of CCAP neurons, CCAP>GFP intensity, and nuclear pMad intensity were selected as representative images.

## Acknowledgments

Research reported in this publication was supported by the National Institute of General Medical Sciences of the National Institutes of Health under award number R01GM068118, and by a Brown University Emerging Area of New Science DEANS Award.

Resources used in this publication were obtained from: DGRC, supported by NIH grant 2P40OD010949; and DSHB, created by created by the NICHD of the NIH and maintained at The University of Iowa. Stocks obtained from the Bloomington Drosophila Stock Center (NIH P40OD018537) were used in this study.

## Author contributions

ENA and KAW designed and analyzed experiments, and revised and approved the manuscript. ENA conducted experiments and drafted the manuscript.

## FOOTNOTES

The abbreviations used are: BMP, bone morphogenetic protein; Gbb, Glass bottom boat; Dpp, Decapentaplegic; Wit, Wishful thinking; Sax, Saxophone; PC, proprotein convertase; R-Smad, receptor-mediated Sma/Mad protein; Tkv, Thickveins; Mad, Mothers against decapentaplegic; S2, Schneider 2; SP, signal peptide; EV, empty vector; GLHT, general linear hypothesis test; PCV, posterior cross vein; CCAP, crustacean cardioactive peptide.

## Footnotes

1 Sue Chien, Mitch Psotka, and Kristi Wharton, personal communication

